# Polymer folding through active processes recreates features of genome organization

**DOI:** 10.1101/2022.12.24.521789

**Authors:** Andriy Goychuk, Deepti Kannan, Arup K. Chakraborty, Mehran Kardar

**Affiliations:** Institute for Medical Engineering and Science, Massachusetts Institute of Technology, Cambridge, MA 02139, United States; Department of Physics, Massachusetts Institute of Technology, Cambridge, MA 02139, United States; Department of Chemical Engineering, Massachusetts Institute of Technology, Cambridge, MA 02139, United States; Ragon Institute of Massachusetts General Hospital, Massachusetts Institute of Technology and Harvard University, Cambridge, MA 02139, United States; Department of Chemistry, Massachusetts Institute of Technology, Cambridge, MA 02139, United States

**Keywords:** Polymer Mechanics, Genome Organization, Active Processes, Stochastic Processes

## Abstract

From proteins to chromosomes, polymers fold into specific conformations that control their biological function. Polymer folding has long been studied with equilibrium thermodynamics, yet intracellular organization and regulation involve energy-consuming, active processes. Signatures of activity have been measured in the context of chromatin motion, which shows spatial correlations and enhanced subdiffusion only in the presence of adenosine triphosphate (ATP). Moreover, chromatin motion varies with genomic coordinate, pointing towards a heterogeneous pattern of active processes along the sequence. How do such patterns of activity affect the conformation of a polymer such as chromatin? We address this question by combining analytical theory and simulations to study a polymer subjected to sequence-dependent correlated active forces. Our analysis shows that a local increase in activity (larger active forces) can cause the polymer backbone to bend and expand, while less active segments straighten out and condense. Our simulations further predict that modest activity differences can drive compartmentalization of the polymer consistent with the patterns observed in chromosome conformation capture experiments. Moreover, segments of the polymer that show correlated active (sub)diffusion attract each other through effective long-ranged harmonic interactions, whereas anticorrelations lead to effective repulsions. Thus, our theory offers non-equilibrium mechanisms for forming genomic compartments, which cannot be distinguished from affinity-based folding using structural data alone. As a first step toward disentangling active and passive mechanisms of folding, we discuss a data-driven approach to discern if and how active processes affect genome organization.

---

The folding of various biopolymers into specific conformations is vital for cellular function. Decades of research on equilibrium polymer theory have revealed basic principles of sequence-controlled folding (1–6). Specifically, polymers composed of sequences of chemically distinct monomers can, via affinity-based monomer-monomer and monomer-solvent interactions, fold into particular shapes despite the dramatic loss of conformational entropy. These ideas range from simple hydrophobic effects that explain the positioning of residues within globular proteins (7–9), to complex free energy landscapes that precisely predict protein structure (1, 5). More recently, analogous concepts have been applied to the folding of interphase chromatin—a complex heteropolymer consisting of genomic DNA and associated proteins (10).

Advances in imaging and sequencing technology have revealed that transcriptionally active euchromatin, which typically resides in the nuclear interior, spatially segregates from transcriptionally silent heterochromatin at the nuclear periphery (11). Chromosome conformation capture (3C) experiments measure this segregation via an enrichment of contact frequencies within euchromatic (A) and heterochromatic (B) regions and depletion of contacts between them (12). These “A/B-compartments” are thought to form because of pairwise attraction between chromatin segments with similar histone modifications, which leads to equilibrium microphase separation (13). Borrowing thermodynamic ideas from protein folding, increasingly sophisticated equilibrium models have designed interaction landscapes to simulate genomic structures that recapitulate 3C data (14–22). This success is remarkable, yet somewhat surprising, since the intracellular environment is far from equilibrium.

Many chromatin-associated proteins are enzymes (23–25) that break detailed balance by turning over chemical energy (26), such as ATP or metabolites, to perform nonequilibrium reactions and/or exert mechanical forces (27), cf. Fig. 1. Such active processes characteristically lead to faster motion that cannot be explained by thermal fluctuations alone (28). Indeed, the subdiffusion of chromosomal loci slows down in the absence of ATP in both bacteria and eukaryotes (29). In addition, nucleus-wide tracking of fluorescently labeled histones has unveiled micron-scale regions of correlated chromatin motion, which disappear after ATP depletion or ATPase inhibition (30, 31). These active processes also vary along the genome, thereby driving heterogeneous motion (32). The apparent diffusion coefficient of individual loci depends on their location in sequence space (33) and on transcriptional state (34), although the precise effect of transcription is debated (35, 36). Similarly, histone tracking experiments have measured faster motion in active euchromatin in the nuclear interior than in heterochromatin at the nuclear periphery (36, 37). These observations point towards the presence of sequence-controlled active forces that affect the polymeric genome’s mobility. To study how such active processes may contribute to shaping genome structure, we need a theory that can link active polymer dynamics to folding patterns (38).

**Fig. 1.**
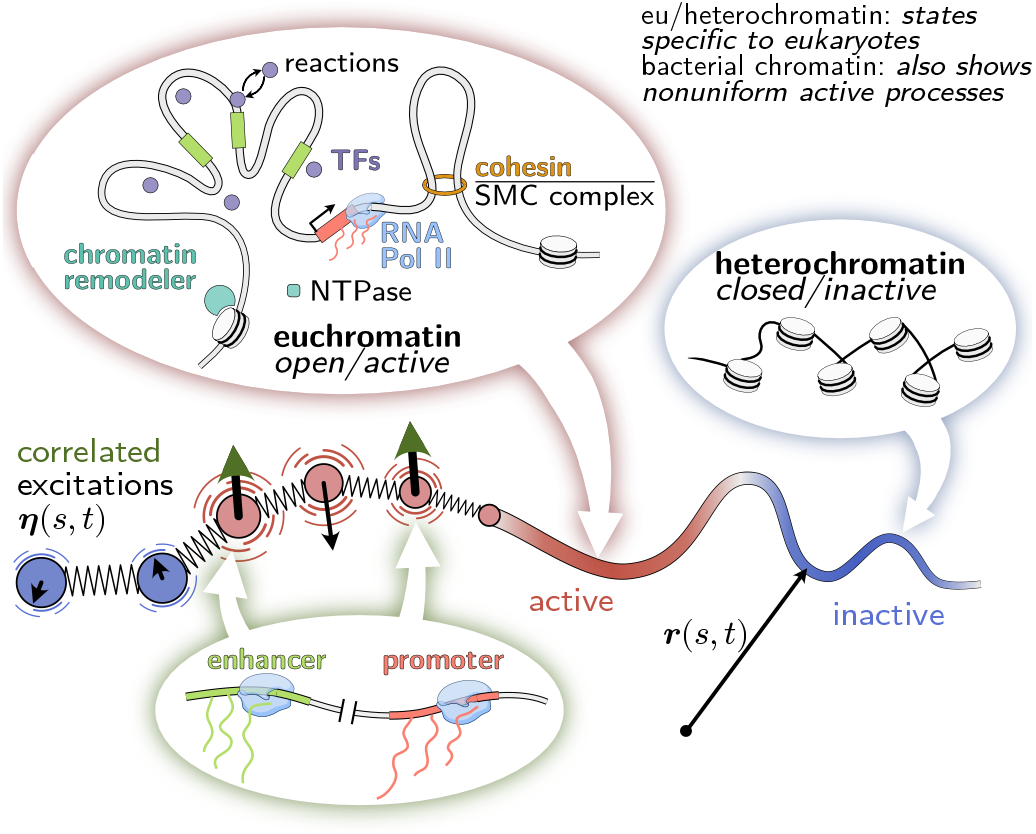
Model of the genome as an active heteropolymer. The genome acts as a scaffold for a myriad of active processes. Motor proteins, such as RNA polymerase II and loop extruding factors, consume chemical energy to exert forces on DNA (24). These nucleoside triphosphatases (NTPases) hydrolyze nucleoside triphosphates (NTP) such as ATP, thereby switching to an NDP-bound state, and restore their NTP-bound conformation by nucleotide exchange (76). For example, specialized proteins actively modify DNA-bound histones and transcription factors via nonequilibrium reactions (23, 77, 78). In the context of eukaryotes, active processes are enriched in transcriptionally active euchromatin compared to inactive heterochromatin. In our coarse-grained model, we describe active processes by random forces that have larger magnitude and thus drive faster diffusion in active regions than in inactive regions. We also stipulate that the active forces at different loci are not necessarily independent. We hypothesize that correlated excitations could arise, for example, from the coordinated activation of an enhancer and its cognate promoter.

By modeling activity via persistent monomer motion, past work has predicted nonequilibrium phenomena such as coherent motion and polymer collapse or swelling (39–49). However, these studies consider uniform activity along the polymer and thus cannot explain heterogeneous folding patterns. More recent models have incorporated non-uniform activity via active forces that vary in magnitude along the chain (50–55), akin to a local effective temperature (56). Simulations have shown that large (30-fold) activity differences can drive phase separation between different polymer regions (50), analogous to mixtures of active and passive particles (57, 58), although smaller activity differences are sufficient in polymeric mixtures (59). Finer grained models have incorporated motor activity via force dipoles (60–62) that align tangentially with the chain (63–66), or through explicit simulation of translocating proteins (67). However, an analytic framework that explains why and how non-uniform activity can fold polymers is still lacking. Moreover, prior work has assumed that active processes at distinct genomic loci are statistically independent. In the context of chromatin, however, we hypothesize that correlations could arise from the coordinated transcriptional activation of different regions, such as enhancers and corresponding promoters (68, 69) or coregulation of genes by common transcription factors (70–72).

To address these open issues, we study a model of a polymer that is driven by correlated active forces with non-uniform magnitude. Our continuum theory shows that active (A) regions of the polymer expand and bend, whereas inactive (B) regions contract and straighten out. Therefore, increased activity within euchromatin could help preserve its expanded state and increase its accessibility to active proteins. Using polymer simulations, we find that even modest activity ratios (two to ten-fold) can recapitulate the degree of A/B compartmentalization observed in Hi-C data. Moreover, we find that distinct loci experiencing correlated active forces will effectively attract, while anticorrelations lead to repulsion. Our results provide a nonequilibrium mechanism that links activity-driven correlated motion to the folding patterns observed in Hi-C data. Finally, we derive an analytical mapping from our active polymer model to an effective equilibrium model where folding is determined by pairwise affinities. These two models are indistinguishable based on structural data alone, raising the need for future dynamic measurements. For example, our model assumptions could be tested via measurements of pairwise velocity correlations of specific loci. Furthermore, given ensemble-averaged conformational data of a polymer, our analytical theory enables us to propose an activity profile that could reproduce the observed steady state. By comparing the inferred activity profile to DNA-binding patterns of chromatin-associating proteins, one could determine whether active processes contribute significantly to certain folding patterns. Taken together, our results provide a new avenue for analyzing and interpreting data on chromosomes, and have broad implications for active polymer systems.

## Model

To study the folding of an active polymer, we combine analytical calculations (of a linear, continuous model) and Brownian dynamics simulations (of a discrete chain). For our theoretical analysis, we idealize the polymer as a space curve ***r***(*s, t*), where *s* is a continuous, dimensionless material coordinate along the polymer backbone. The large-scale dynamics of a polymer are well-described by the Rouse model, an extensile chain of material points that interact through Hookean springs *κ* (39, 73–75),

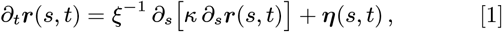

where *ξ* is the drag friction with the surrounding solution and ***η***(*s, t*) is a zero-mean Gaussian random displacement velocity field which we refer to as “excitations”. In general, the covariance of the excitations at different material points is given by

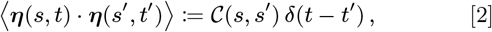

on timescales longer than the decorrelation time. For a passive polymer that reaches thermal equilibrium, the excitations are determined by the heat bath, 𝒞 (*s, s*′) = 6*k*_*B*_ *Tξ*^−1^ *δ*(*s* − *s*′), which follows from the fluctuation-dissipation theorem. However, sequence-specific active processes that stir the polymer through random forces will drive the system away from thermal equilibrium. These “athermal excitations” may vary in magnitude along the polymer and exhibit sequence-specific correlations,

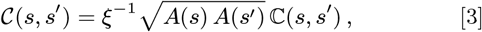

where *A*(*s*) is the activity at locus *s* (in units of dissipated energy), and ℂ(*s, s*′) is the “normalized” correlation function. Figure 1 depicts possible molecular drivers of these athermal excitations in the context of the genome. Note that Eq. (3) is directly proportional to the pairwise velocity correlation function (Supporting Information 1 C). Thus, by describing the polymer response to these excitations, our model links patterns of correlated motion with patterns of folding.

### Brownian dynamics simulations of a discrete chain

To numerically test our theoretical analysis, we developed Brownian dynamics simulations of a discretized Rouse chain, Eq. (1), where each of the *n* ∈ [1, *N*] monomers represents a Kuhn segment with characteristic length *b*. In our simulations, we use the Kuhn length *b* and the average diffusion coefficient *D*_0_ of a free monomer, which is related to the average activity *A*_0_ = ∑_*n*_ *A*_*n*_*/N* via *D*_0_ = *A*_0_*/*(6*ξ*), to define the stiffness of the springs connecting neighboring monomers, *κ/ξ* = 3*D*_0_*/b*^2^. Having discretized all fields of our continuum theory, the covariance matrix between the athermal excitations of different beads is then given by

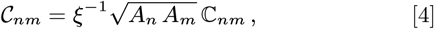

where we have defined the Pearson correlation matrix ℂ_*nm*_ ∈ [−1, 1]. To implement more realistic polymer simulations, including self-avoidance and a hard confinement, we adapt the “polychrom” software package as described in the Methods. These simulations allow us to test the continued validity of results from our theory in a strongly nonlinear setting that is inaccessible to analytical calculations.

### Steady-state polymer conformation

We analytically solve the linearized active polymer model [Eq. (1)] via a Rouse mode decomposition (79), where 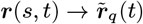 and 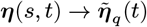 indicate Fourier transforms. For a compact notation, we concatenate all Rouse modes row-wise into a matrix ***R***(*t*) with rows 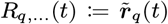. Analogously, we define the random velocity mode matrix ***H***(*t*) with rows 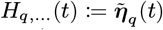, which has zero mean and covariance ⟨***H***(*t*) · ***H***^†^(*t*′) := ***C*** *δ*(*t* − *t*′)⟩. In the resulting Rouse mode dynamics,

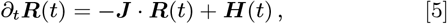

the response matrix ***J*** encodes all material properties (Methods) and is diagonal for a homogeneous Rouse polymer [Eq. (1)], *J*_*qk*_ = *ξ*^−1^*κq*^2^ *δ*_*qk*_. In contrast, athermal excitations that break translational invariance are characterized by off-diagonal entries in their covariance matrix 𝒞 (*s, s*′) → *C*_*qk*_. Although this coupling precludes an isolated analysis of individual Rouse modes, one can derive an exact expression for the long time limit of the second Rouse moment (Methods) as

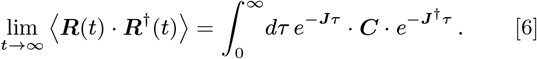

Note, however, that the above equation describes polymer confirmations on average, which are liquid-like in the sense that ⟨***R***(*t*) · ***R***^†^(*t*) ⟩ ≫ ⟨***R***(*t*) ⟩ · ⟨***R***^†^(*t*) ⟩. The resulting “folding” is thus distinct from most proteins, which form globules with a well defined conformation, ⟨***R***(*t*) ⟩ · ⟨***R***^†^(*t*) ⟩ ∼ ⟨***R***(*t*) · ***R***^†^(*t*) ⟩. Nevertheless, our analysis reveals how inhomogeneous excitations alone can effectively give rise to *patterned* polymer conformations by coupling different mechanical modes. To quantify these patterns, we transform Eq. (6) back into real space, which yields the spatial correlation between pairs of material points. Subsequently, we determine their pairwise mean squared separation and tangent autocorrelations.

## Results

### Local activity modulations induce long-range correlations akin to bending

To elucidate how active processes affect a polymer’s conformation, we first study a minimal scenario with inhomogeneous activity *A*(*s*) represented by *statistically independent* excitations 𝒞 (*s, s*′) = *ξ*^−1^ *A*(*s*) *δ*(*s* − *s*′). Simulations have shown that less active monomers localize closer to the boundary of a hard confinement than their more active counterparts (50, 51). This positioning trend reverses if the volume packing fraction is small or if the confinement is soft (51), and can be forcibly inverted by introducing selective monomer–boundary interactions (50), or self-attraction between inactive monomers (55). Thus, past theoretical work has shown that activity differences can reproduce nuclear positioning of (active) euchromatin and (inactive) heterochromatin. However, it is not yet clear how and why active processes affect *polymer shape*, a question that we now address.

Equation (6) predicts the preferred polymer conformation in response to any predetermined profile of activity [see Supporting Information 2 A.3 for Green’s function kernels]. As an example, we focus on sinusoidal activity modulations *A*(*s*) = *A*_0_ [1 + *ϵ* cos(*s/λ*)] around an average value *A*_0_, with amplitude *ϵA*_0_ and characteristic length *λ*. These excitations elicit a spatial correlation between pairs of material points,

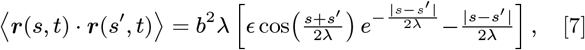

which we use to calculate the pairwise mean squared separation in terms of the Kuhn length 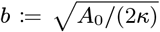 [below the diagonal in Fig. 2B]. In comparison to a reference polymer with uniform activity (*ϵ* = 0), we observe that active polymer segments locally expand while inactive segments locally condense [above the diagonal in Fig. 2B], resembling the morphology of euchromatin and heterochromatin (81, 82). Thus, our theory suggests that active processes like transcription could lead to chromatin decondensation, which might form a positive feeback loop by further increasing genome accessibility to the transcription machinery (83). Indeed, it was shown experimentally that euchromatin requires ATP and thus dissipation of energy to preserve its expanded state (30).

**Fig. 2.**
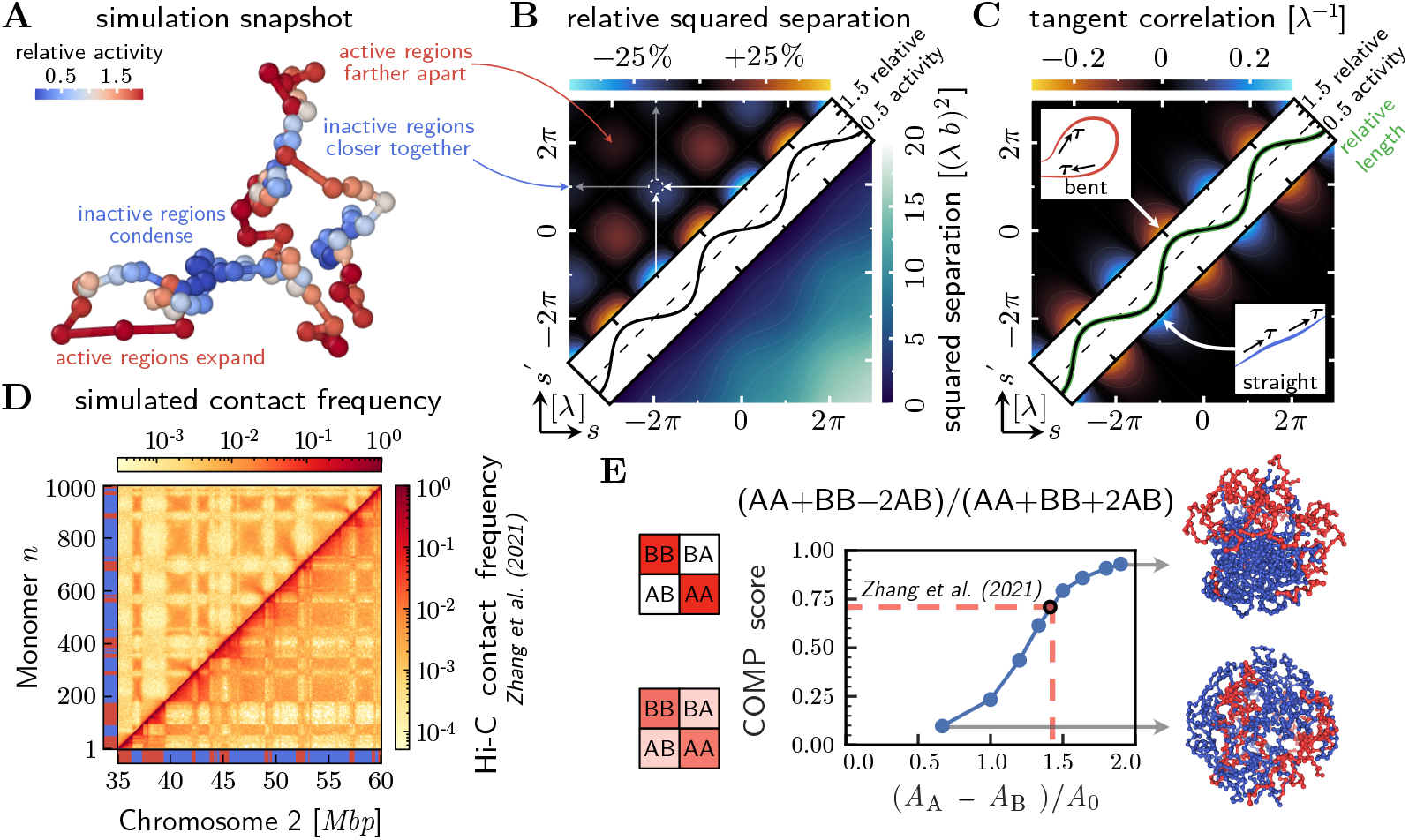
Activity modulations induce polymer folding. **A)** Simulation snapshot of a Rouse chain composed of 100 monomers, driven by statistically independent athermal excitations that vary in magnitude, with local activity *A*(*s*) = *A*_0_ [1 + *ϵ* cos(*s/λ*)] and *ϵ* = 0.5. We plot the same activity profile normalized by its average value, *A*(*s*)*/A*_0_, along the diagonal of panels B and C. **B)** *Below the diagonal:* Mean squared separation between pairs of material points with coordinates *s* and *s*′ along the polymer backbone. *Above the diagonal:* Change in mean squared separation relative to a Rouse polymer with homogeneous activity *A*_0_. Active regions locally expand while inactive regions locally condense. Distant active polymer segments get farther apart while inactive polymer segments come closer together (dashed circle). **C)** Correlation between tangent vectors at different material points. A local increase in activity leads to polymer bending (anti-correlations), while a local decrease in activity leads to polymer straightening (positive correlations). Local expansion and contraction of the polymer backbone are reflected in the relative contour length (plotted on the diagonal), which is proportional to the relative level of activity. **D)** *Below the diagonal:* Hi-C data from Ref. (80) showing the contact frequencies between pairs of genomic coordinates within a section of Chromosome 2 in murine erythroblast cells. *Above the diagonal:* Simulated contact probability between pairs of monomers of a 1000-mer self-avoiding chain in a hard spherical confinement. The activity difference between active (A, red) and inactive (B, blue) monomers is chosen to match the degree of compartmentalization observed in the Hi-C data. **E)** Simulated contact frequency maps show increasing compartmentalization (COMP score) as a function of the normalized activity difference between A and B regions. Dashed lines illustrate the activity difference (*A*_*A*_*/A*_*B*_ ≈ 6) required to reproduce the COMP score observed in the Hi-C data of Ref. (80). Representative simulation snapshots show phase separation within the polymer.

Consistent with prior work (50, 52, 54, 55), we also find that pairs of active segments get farther apart while inactive segments come closer together [above the diagonal in Fig. 2B]. To investigate the shape changes associated with this segregation, we measure the pairwise alignment of tangent vectors, ***τ*** (*s, t*) := ∂_*s*_***r***(*s, t*), at different material points (tangent autocorrelation),

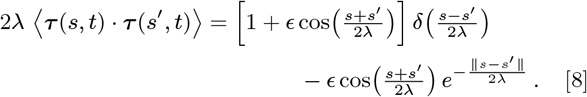

The last term clearly demonstrates that local activity modulations induce correlations (effective pairwise couplings) between distant material points. Specifically, Figure 2C shows that increased activity within a segment of size *λ* effectively induces bending (anti-correlations between tangent vectors), while decreased activity leads to straightening (positive correlations between tangent vectors). Heuristically, active monomers “run away” from their inactive neighbors, effectively bringing these neighbors together into a loop-like conformation. Thus, we conclude that segmented activity variations lead to the emergence of spontaneous curvature.

The first term in Eq. (8) describes local compaction (or expansion) of the polymer backbone from reduced (or increased) activity. Similar changes in the local contour length could be induced by variations in tension, i.e. the spring constant *κ*(*s*). Therefore, it is natural to inquire if conformational changes due to activity variations can be equivalently achieved in a system in thermal equilibrium through variations in the local mechanical properties of the polymer. However, as shown in the Supporting Information 2 A.2, modulations in tension fail to produce any long-ranged tangent correlations akin to polymer bending. Changes in the spring stiffness only affect the distance between material points (bond length) and have no effect on the bond angles. Similarly, variations of the drag friction coefficient *ξ*(*s*), for example, due to differences in monomer size, do not change the polymer conformation (Supporting Information 2 A.1). This is because in equilibrium with a thermal heat bath, polymer conformations sample a Boltzmann distribution determined by a free energy, independent of dynamic effects such as friction. Thus, in the absence of long-ranged interactions, activity differences, which can act globally, are required to fold a Rouse polymer into specific conformations.

### Activity differences recapitulate A/B-compartments in simulated contact maps

We next apply our model to study the formation of A/B-compartments, a cornerstone of eukaryotic genome organization (12). To make a meaningful comparison to Hi-C data, we model a region of Chromosome 2 in murine erythroblast cells (80) as an active, self-avoiding polymer in a nuclear confinement (Methods), where each monomer corresponds to 25 kbp. We ask whether the compartmentalization observed in Ref. (80) [Fig. 2D, below the diagonal] can be reproduced purely by differences in the magnitude of athermal excitations in active (A) and inactive (B) regions. To that end, we derive the identities of A and B monomers from the data in Ref. (80), as discussed in the Methods. The simulated contact frequencies between pairs of monomers display a checkerboard pattern featuring strong B–B contacts, despite the lack of explicit attractive interactions between monomers [Fig. 2D, above the diagonal]. While A–A contacts in our simulations are weaker than in the data of Ref. (80), this could be remedied by including weak self-attraction among all monomers (13) or, as discussed in the next section, by introducing correlated excitations. Combining active and passive mechanisms of compartmentalization will invariably fit the data better (55), but the existence of A/B-compartments in our minimal model demonstrates that activity differences are capable of contributing to genome organization.

To explore how the degree of compartmentalization varies with the activity difference between A and B regions, we extract a scalar order parameter (“compartment score”, COMP) from both simulated and experimental contact maps (Methods). This compartment score, defined in Ref. (84), measures the contrast in the checkerboard pattern as the normalized contact frequency difference between same-type and different-type chromatin, namely COMP = (AA + BB − 2AB)*/*(AA + BB+2AB). We find that the compartment score increases in a sigmoidal fashion with the activity difference between A and B regions [Fig. 2E], indicating a typical scale for onset of compartmentalization in our simulations. This activity difference scale depends on many parameters, including the A and B block sizes (59) and the capture radius used to construct the contact frequency map. In this particular example, we find nontrivial compartmentalization for activity differences as small as the average level of activity, *A*_A_ − *A*_B_ = *A*_0_ = (*A*_A_ + *A*_B_)*/*2. Note that in our analytic theory, which describes a phantom chain without volume exclusion, compartmentalization, as detected in the mean squared separation, is simply a linear response to the activity difference, Eq. (7).

Finally, the compartment score curve in Fig. 2E can be used to read off the activity difference required to reproduce A/B compartmentalization in a given Hi-C dataset. While the degree of chromatin compartmentalization will vary by cell type, a whole genome analysis of the murine erythroblast data of Ref. (80) [Fig. 2E] suggests a score of COMP ∼ 0.71, which corresponds to an activity ratio of *A*_A_*/A*_B_ ≈ 6. We used this inferred activity ratio in the polymer simulations depicted above the diagonal in Fig. 2D.

While the activity ratio cannot be measured directly, one can use the monomer mean squared displacement, MSD(*t*) = *D*^app^ *t*^*α*^, as a proxy. On sufficiently short time scales in a phantom Rouse chain, the ratio of anomalous diffusion coefficients in active and inactive regions is identically the activity ratio,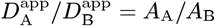. However, the value of *D*_app_ and *α* in active and inactive regions will depend on the observation time window and the microscopic properties of the chain, as shown in our nonlinear simulations (Supporting Information 3 C). Thus, our predicted activity ratio serves as an upper bound for the ratio of MSDs in A and B regions which can be extracted from measurements of euchromatic and heterochromatic motion (36).

### Correlated active processes create compartments

Experiments that track the movement of GFP-tagged histones have demonstrated spatial correlations in chromatin motion that depend on RNA polymerase II activity and on ATP (30, 31). While these experiments cannot relate spatial and genomic proximity, Ref. (85) has observed pairwise correlated movement of specific loci. Motivated by these experiments, we hypothesize that correlated movement could be driven by correlated active forces, which produce athermal excitations with a non-diagonal covariance 𝒞 (*s, s*′). This hypothesis is plausible if the active processes at distinct genomic locations are not completely independent.

To model sequence correlations, we define a heteropolymer with three types of monomers (+, −, and neutral). We then introduce a stochastic process that has opposing effects on + and − monomers, but does not affect neutral monomers. The choice of + and − as labels for the monomer type evokes the analogy of a charged heteropolymer (for example, a polyelectrolyte or polyampholyte) in a fluctuating electric field. In this example, the random electrical forces at a given time will point in the same direction for monomers of the same charge, but in opposite directions for monomers of opposing charge. Here, charge can be interpreted as any biochemical signature of a monomer (such as the methylation status of a chromosomal locus) that mediates a selective response to a long-ranged active control process.

Based on these ideas, we construct a sample Pearson correlation matrix for the excitations of a polymer with an alternating pattern of +, −, and neutral monomers [Fig. 3A]. Using this Pearson correlation matrix as input and assuming a homogeneous level of activity, see Eq. (4), we then perform Brownian dynamics simulations featuring self-avoidance and a spherical confinement. Figure 3B shows, below the diagonal, that contacts between loci driven by correlated excitations are enhanced, whereas contacts between anti-correlated loci are depleted. These folding patterns are accompanied by long-range correlations between the tangent vectors at distant material points, as shown in our theoretical calculations (Supporting Information 2 A.6). In addition to the pattern of correlations, we superimpose activity modulations, *A*_*n*_, such that charged (±) monomers are active and neutral monomers are inactive. We find an enhancement of contacts between inactive regions, leading to a full checkerboard pattern [Fig. 3B, above the diagonal]. Thus, correlated active forces and activity differences offer complementary, non-equilibrium mechanisms for the formation of genomic compartments.

**Fig. 3.**
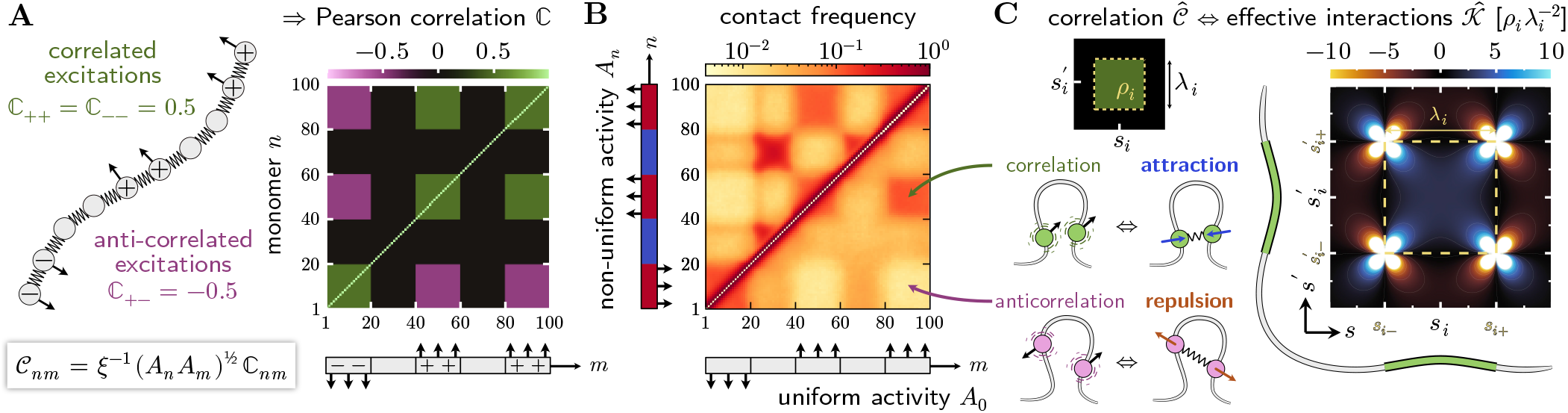
Correlated activity leads to compartments via an effective attraction. **A)** Pearson correlation matrix that relates the directions of the random forces acting on different monomers. We construct this matrix by assigning each monomer a chemical identity (+, −, or neutral) whereby monomers of the same chemical species experience correlated excitations, while monomers of different species experience anti-correlated excitations. In the schematic below the matrix, up arrows indicate + monomers and down arrows indicate − monomers. **B)** Simulated contact frequencies between pairs of monomers of a 100-mer self-avoiding chain in a spherical confinement. The polymer is driven by athermal excitations with the Pearson correlation matrix shown in panel A. *Below the diagonal:* When activity is uniform, contacts between monomers that experience correlated excitations are enhanced, while anticorrelated excitations lead to a depletion of contacts. *Above the diagonal:* We introduce activity modulations where + and − monomers have activity 1.9*A*_0_ (red), and neutral monomers have activity 0.1*A*_0_ (blue). As in Fig. 2, inactive regions form compartments, without disrupting the folding pattern caused by correlated excitations. **C)** Correlated excitations fold an active polymer analogously to effective harmonic interactions in a passive polymer, cf. Eq. (14). To illustrate these couplings, we consider two polymer segments of equal length *λ*_*i*_ located at coordinates *s*_*i*_ and 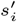, which experience excitations with correlation coefficient *ρ*_*i*_ (green box). Positive correlations (*ρ*_*i*_ > 0) lead to an effective attraction (blue) between these segments through weak, long-ranged harmonic interactions. Conversely, anti-correlated excitations (*ρ*_*i*_ < 0) lead to an effective repulsion (orange) between these segments. If the coordinates *s*_*i*_ and 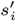 coincide (that is, if we consider correlated excitations within a contiguous segment), then the correlation coefficient is positive and the segment condenses.

### The effects of correlated, non-equilibrium excitations on polymer shape can be recapitulated by an equilibrium model with intersegment interactions

The linear active polymer, driven by athermal Gaussian excitations with covariance 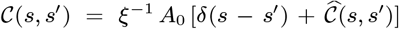, maintains a Gaussian steady-state conformation. The Gaussian steady state is fully characterized by its two-point correlation function ⟨***r***(*s, t*) · ***r***(*s*′, *t*) ⟩, which can be computed in terms of the prescribed entries of 𝒞 (*s, s*′). To illuminate the characteristics of this *active folding*, we note that any Gaussian steady state can be regarded as the thermal equilibrium weight of a phantom polymer with additional Hookean springs 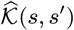 that har-monically couple pairs of material points (18). After setting a temperature scale by *A*_0_ = 6*k*_*B*_ *T*, one can obtain the harmonic couplings by inverting the two-point correlation function ⟨***r***(*s, t*) · ***r***(*s*′, *t*)⟩, and as shown in the Methods,

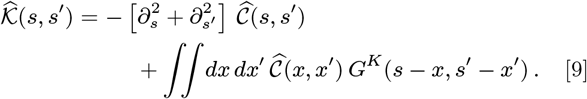

The Green’s kernel has the most convenient representation in polar coordinates *G*^*K*^(*α* cos *ϕ, α* sin *ϕ*) = − 6 cos(4*ϕ*)*/*(*πα*^4^). We note that, while the active polymer folds due to constraints on the excitations (i.e. correlations), in the corresponding passive polymer constraints are introduced in the form of springs. These harmonic couplings can be positive or negative, mediating pairwise attraction or repulsion between distant material points. The first term of Eq. (9) indicates that pairs of polymer segments experiencing maximally correlated excitations will show effective pairwise attraction, while maximally anticorrelated excitations lead to repulsion. By using a box-shaped correlation function to study the second term of Eq. (9), we confirm that correlated excitations *in general* induce pairwise attraction [Fig. 3C], whereas anti-correlated excitations lead to repulsion.

These results can be heuristically understood via the dynamics of the end-to-end distance vector of, say, a trimer (*N* = 3). For such a trimer, fluctuations of the end-to-end distance Δ are driven by a noise term with variance ⟨[***η***Δ(*t*)]^2^⟩ = ⟨[***η***_1_(*t*) − ***η***_3_(*t*)]^2^⟩ = ⟨[***η***_1_(*t*)]^2^⟩ + ⟨[***η***_3_(*t*)] ⟩ − 2⟨***η***_1_(*t*) · ***η***_3_(*t*)⟩.

When the excitations are independent and equal in magnitude, one has ⟨ [***η***Δ(*t*)]^2^ ⟩ = 2 ⟨ [***η***_1_(*t*)]^2^ ⟩. In comparison, anticorrelated excitations (⟨***η***_1_(*t*) · ***η***_3_(*t*) ⟩ < 0) increase the variance of the end-to-end separation, whereas correlated excitations (⟨***η***_1_(*t*) · ***η***_3_(*t*) ⟩ > 0) cause the end points to come closer together. While these effects are diminished for longer polymers, they provide basic insights into the effective harmonic interactions that lead to the contact frequency map shown in Fig. 3B.

The existence of an analytical mapping between active and passive mechanisms of polymer folding suggests that many equivalent models could explain structural data on chromosomes. In this context, our passive linear model can be regarded as a harmonic approximation to a contact energy landscape, which has been exploited by past theoretical work on genome organization (14–17, 19–21). Our results explain the somewhat surprising success of these equilibrium models in reproducing Hi-C data despite the undeniable presence of active processes (86). As such, how can one experimentally disentangle equilibrium and non-equilibrium mechanisms of compartment formation? Based on static snapshots, our theory can be used to propose a candidate pattern of active processes that drive chromatin towards its observed steadystate conformation. Then, one can test if the inferred activity profile matches orthogonal experimental measurements, such as the DNA-binding patterns of active enzymes, or if passive folding mechanisms are more plausible.

### Inferring an activity profile from an ensemble-averaged polymer conformation

In the following, we infer a candidate map of athermal excitations that could fold a polymer into a desired (target) conformation. To that end, we use our linear theory to predict the activity of each monomer given the mean squared separation between all pairs of monomers [cf. workflow depicted in Fig. 4A]. We test our approach on artificial target conformations, which we generate via simulations of (nonlinear) polymers driven by activity modulations, with profiles corresponding to Fig. 2D and E.

**Fig. 4.**
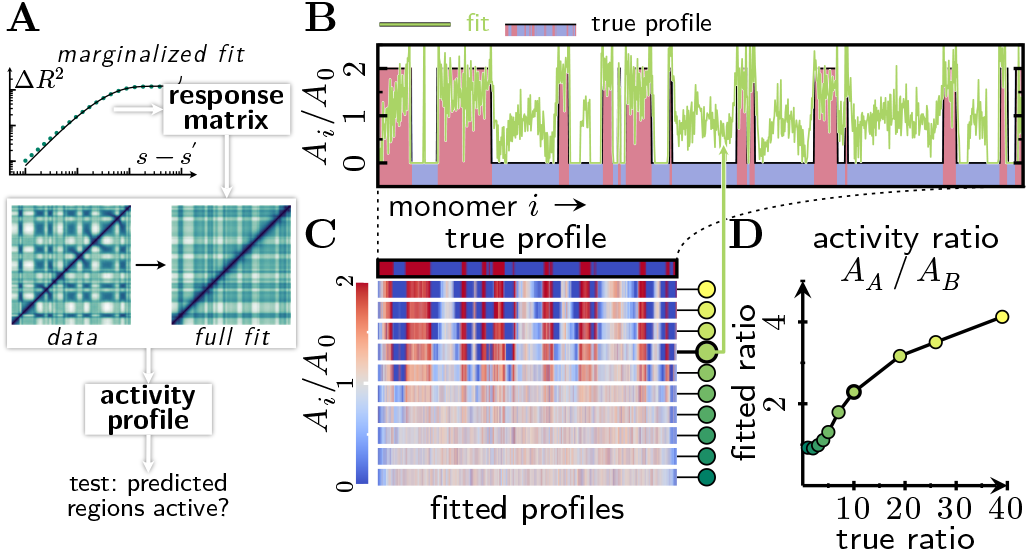
Inferring an activity profile based on structural data. **A)** Data analysis workflow, which takes as input a heatmap with the pairwise mean squared separation between all monomers. First, we infer the mechanical properties of the homogeneous polymer on top of which activity differences will be imposed. We then propose a profile of activity that minimizes the mean squared error between the predicted mean squared separation map and the data. This workflow can be used to identify candidate active regions of the polymer, which could then be tested against alternative measurements. **B)** We applied the workflow shown in A to simulated data on self-avoiding polymers driven by activity modulations in a spherical confinement [cf. Fig. 2E]. We find, in general, good agreement between the inferred activity profile and the true profile used in the simulations. **C)** The fitted activity profiles correctly identify the boundaries between active and inactive regions in the true activity profile. Color-coded circles next to the activity profiles correspond to the plot points (fitted activity ratios) in panel D; highlighted circle indicates activity profile in panel B. **D)** The fitted activity ratio between active (A) and inactive (B) regions correlates with the activity ratios of the athermal excitations used in our simulations. In our linear model, a systematically lower activity ratio is sufficient to reproduce the folding patterns observed in the non-linear simulations.

As it is not clear how nonlinear constraints such as self-avoidance or a hard confinement translate into our linearized theory, we invoke no prior knowledge on the mechanical properties of the polymer. Instead, we adopt a data-driven approach and first determine an effective response matrix ***J*** that approximates the mechanical properties of the simulated polymer (Supporting Information 4 A, 4 B). Next, we set up a numerical optimization scheme whereby we seek the activity profile [Fig. 4B] that minimizes the squared deviation between the mean squared separation map predicted by our theory and our artificial data (Supporting Information 4 C). Inference from linear theory captures the overall block-like structure of the mean squared separation map, but does not account for some of the finer qualitative features observed in the nonlinear simulations (Supporting Information 4 C). We hypothesize that these fitting results could be improved in future studies by introducing constraints on the analytical theory, or by considering excitations that have an effective correlation length in addition to activity modulations.

Correspondingly, the linear model successfully infers the structure of the activity profile used in our nonlinear simulations [Fig. 4B,C]. The inferred and original activity profiles sually appear similar and have comparable amplitudes *A*_*A*_−*A*_*B*_ [Fig. 4B]. However, the inferred ratio of activity *A*_*A*_ */A*_*B*_ is systematically lower than in our simulations [Fig. 4D]. These results suggest that the lack of constraints in the linearized model makes it easier to create folding patterns than in our simulations. Despite these quantitative differences, this approach successfully identifies active and inactive polymer segments in all simulations that show pronounced folding patterns [Fig. 4C].

In closing, we propose a novel approach for the analysis of experimental data on chromatin. As a first test for plausibility, one could compare inferred activity profiles against orthogonal experimental measurements such as ChIP-Seq for ATPases or histone marks associated with active chromatin. After using our theory to find candidate segments with high predicted activity, it would be interesting to measure the effect of biochemical interventions that locally knock out energy-dissipating processes. By testing model predictions against experiments, one can then distinguish if a certain folding pattern is dominated by active processes or passive effects. Finally, the linear theory can be generalized to include both active processes and passive interactions. One could then test the signatures of this hybrid model by measuring the fluctuation dynamics of specific polymer segments (87).

## Discussion

Using our nonequilibrium polymer theory, we have demonstrated that differences in activity and correlations between athermal excitations at different loci can fold a polymer into specific shapes. These liquid-like conformations have a large variability, and are therefore only realized as a population average. Our model could thus be applicable to chromatin, which shows a much larger cell-to-cell variability than stable protein structures (88, 89). In this context, we zoom out to large length scales where chromatin behaves like a Rouse polymer (90–92), and molecular drivers of active processes [Fig. 1] produce effective athermal excitations. Our model then makes several testable predictions that could be further investigated with experiments and theory.

We predict that a local increase in activity should lead to chromatin decondensation [Fig. 2A,B]. The resulting increase in chromatin accessibility could further increase transcriptional activity (83), forming a nonlinear positive feedback loop. Such a feedback loop would, over time, stabilize an open chromatin structure in active regions. Furthermore, our model shows that a local decrease of activity leads to straightening of the polymer backbone, whereas a local increase of activity induces bending [Fig. 2C]. These results could explain an observation in recent simulations, which demonstrated that forces exerted by bound molecular motors can bend polymers into hairpins (67). Such zipped structures also arise in Hi-C maps (93–95) and in our simulated contact maps as jet-like, anti-diagonal features originating from small A regions and extending into the neighboring B compartment [Fig. 2D]. Experiments suggest that these structures could be formed by active loop extrusion (93), which is not explicitly accounted for in our model. Nevertheless, it is interesting that the *spontaneous* loops induced by a local hot spot of activity could still create jet-like features.

In addition to locally deforming the polymer backbone, activity differences lead to a “checkerboard” pattern indicative of compartmentalization. We show that the degree of A/B compartmentalization observed in experimental contact maps can be recapitulated in our simulations with modest activity ratios in the two-fold to ten-fold range [Fig. 2E]. To test if these activity ratios are plausible, one could measure the ratio of the anomalous diffusion coefficients of euchromatic (A) and heterochromatic (B) regions, as has been done in histone tracking experiments (36). Since euchromatin has a higher level of transcriptional activity, we may expect that it will exhibit faster (sub)diffusion than heterochromatin (36, 37). However, the effect of transcription on the subdiffusion of individual loci is controversial (34, 35). One way to explain these conflicting results is to note that diffusion is not only proportional to activity, which increases during transcription, but is also inversely proportional to friction, which increases when the transcriptional machinery binds to the promoter. Concomitant with this idea, it was observed that during transcription inhibition, the subdiffusion of DNA-bound histones increases after RNA polymerase II disassociates (36, 37), but decreases if RNA polymerase II remains bound (36). These observations can be reconciled with our theory, which shows that the polymer conformation is independent of friction changes and only responds to activity differences (Supporting Information 2 A.1).

Another test of activity-induced compartmentalization would be to perform Hi-C experiments after knocking out active processes. However, global ATP depletion runs the risk of glassifying the intracellular environment (96). Existing experiments with transcription inhibition show that A/B-compartments remain, but the contrast in the checkerboard pattern decreases (80, 97). It therefore seems plausible that transcription plays some role in compartmentalization, which must be clarified in future studies (98). We hypothesize that activity differences may complement known mechanisms of eu/heterochromatin segregation, including phase separation mediated by linker histone H1 and heterochromatin protein HP1-*α*, as well as association of heterochromatic domains to the nuclear lamina (10).

In addition, whole-nucleus histone tracking has shown that transcription and ATP-dependent processes are required for coherent motion of chromatin (30, 31). The mechanistic origin of correlated motion is under active research (55, 60, 61, 64, 65, 99, 100) and its consequences for chromatin folding are still unknown. Our theory predicts that regions exhibiting correlated motion will, when driven by sequence-specific athermal excitations, form compartments. This raises the question of whether coordinated transcription of enhancers and promoters could lead to contacts, or “micro-compartments”, as has been observed in increasingly high-resolution contact maps (101–103). However, further research is needed to investigate the relationship between coordinated transcriptional programs and correlated motion. To directly test our model assumptions, one could measure pairwise velocity correlations between specific loci, as a function of transcriptional state and sequence coordinate (85). For example, one could measure whether cis regulatory elements show correlated movement, and then test if transcription inhibition at one locus decorrelates this movement and the associated contacts. A similar procedure could be used to test whether the co-localization of co-regulated genes is a consequence of correlated active processes (70, 71, 104). For example, coordinated transcription factor binding events can initiate processes that break detailed balance (105, 106).

Overall, dynamic measurements will play a major role in disentangling active and passive mechanisms of chromatin folding, which we have demonstrated to be indistinguishable based on Hi-C data alone. In future work, we will identify dynamic signatures of activity (87) that can be extracted from trajectories of genomic loci (107) in order to examine the role of active processes in chromatin folding. As a complementary technique, we have here presented a proof of concept that an activity profile can be inferred from structural data on a polymer. In future work, we propose to extend this method to DNA-MERFISH data (108), which measures the pairwise mean squared separation of genetic loci along with other attributes such as transcriptional (108) and epigenetic state (109, 110).

In closing, we note that our theory also has broad implications beyond chromatin folding. We hypothesize that one could, for example, use electromagnetically driven colloidal (Janus) particles to engineer active polymers that fold into desired conformations (111), by applying targeted excitations through dynamic light patterns. Furthermore, on large length scales, our model shares its mathematical structure with mechanical models of membranes. Interestingly, it was recently shown that Min proteins can deform giant unilamellar vesicles by binding to the membrane (112). While it was hypothesized that Min proteins induce spontaneous curvature (113) and that this could even affect their binding kinetics (114), the underlying mechanism remains unknown. Our theory could provide some hints as to how reactions themselves could effectively induce spontaneous curvature in membranes. Overall, our analysis motivates future research into the role that active processes play in determining the conformation of a variety of pattern-forming systems.

## Methods

### Derivation of the steady-state polymer conformation

We now outline a brief derivation of the steady-state polymer conformation, starting from Eq. (5); a more elaborate calculation is provided in the Supporting Information 1 B. Given an unknown matrix-valued function ***H***(*t*), which encodes a possible trajectory of the excitations, Eq. (5) is formally solved by

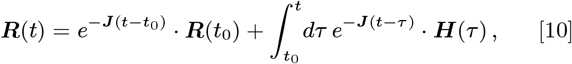

where the first term vanishes in the limit *t* − *t*_0_ → ∞. We use the formal solution for a single trajectory of the Rouse matrix, Eq. (10), to determine the second Rouse moment, ⟨***R***(*t*)·***R***^†^(*t*)⟩, in response to athermal excitations with covariance ⟨***H***(*t*) ***H***^†^(*t*′) ⟩ := ***C*** *δ*(*t* − *t*′). To that end, we multiply Eq. (10) with its conjugate transpose and average over many trajectories.

Both in the limit of late times (*t* → ∞ for arbitrary *t*_0_), or equivalently early reference times (*t*_0_ → − ∞ for arbitrary *t*), one then finds

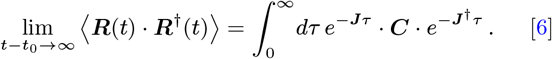

Here and henceforth, we consider reference times that lie infinitely far in the past (*t*_0_ → − ∞), and therefore omit the time limit 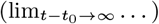 for brevity of notation. For polymers whose material properties such as line tension and friction are homogeneous, the response matrix is diagonal (*J*_*qk*_ ≡ *J*_*qq*_ *δ*_*qk*_) and Eq. (6) evaluates to:

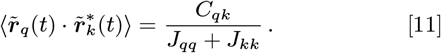

For a Rouse polymer (*J*_*qk*_ = *ξ*^−1^*κq*^2^ *δ*_*qk*_) that, as discussed in the main text, is driven by correlated athermal excitations with mode covariance 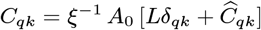, one has:

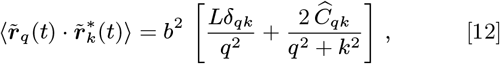

where the characteristic length is given by 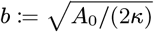. In general, however, the response matrix will be non-diagonal and we need to use a perturbation approach (Supporting Information 1 E) to approximate the matrix exponentials in Eq. (6).

### Steady-state conformation of a passive polymer

We now consider the steady-state conformation of a polymer of length *L*, which is in thermal equilibrium with a heat bath of temperature *T*. The passive polymer is governed by reciprocal interactions and therefore has a Hermitian response matrix, ***J*** = ***J*** ^†^. In thermal equilibrium, the excitations are statistically independent and homogeneous, with covariance 𝒞 (*s, s*′) = *ξ*^−1^*A*_0_ *δ*(*s − s*′) and activity *A*_0_ = 6*k*_*B*_ *T*. Because different excitation modes are now independent, *C*_*qk*_ = *ξ*^−1^*A*_0_ *Lδ*_*qk*_, Eq. (6) evaluates to

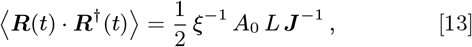

so that polymer folding can only be induced by non-diagonal elements in the response matrix ***J***. We herein assume that the response matrix is dominated by topological connectivity of neighboring material points and by homogeneous mechanical features of the polymer backbone. Therefore, we decompose the response matrix into a dominant diagonal contribution 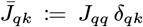 and a weak off-diagonal contribution 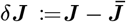, so that

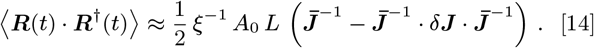

To enforce polymer folding in our equilibrium model, we introduce additional Hookean springs to Eq. (1),

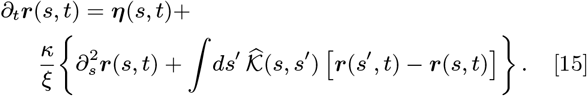

With 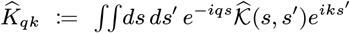 as convention for Fourier transforms, the diagonal and off-diagonal components of the response matrix are given by 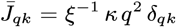 and 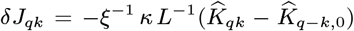, respectively. Thus, Eq. (14) evaluates to:

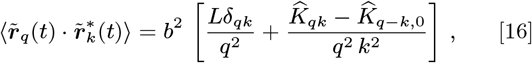

where the characteristic length is given by 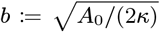, as before. Note that the homogeneous Fourier modes of the harmonic interaction map, *q* = 0 and *k* = 0, cancel out in the polymer’s Langevin equation (15), and thus also in the steadystate conformation, Eq. (16). Therefore, in the following we assume 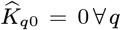. Equating Eq. (16) and Eq. (12), and transforming back into real space, one obtains Eq. (9).

### Rouse polymer simulation details

To test the approximations in our analytical theory and visualize individual polymer conformations, we perform Brownian dynamics of a discretized Rouse chain without self-avoidance or confinement: https://github.com/kannandeepti/active-polymers. Since the average monomer diffusion coefficient *D*_0_ only rescales time and has no effect on the steady state conformation, we arbitrarily set *D*_0_ = 1 and the Kuhn length *b* = 1, such that all simulated length scales are now in units of Kuhn lengths. We integrate the discrete version of Eq. (1) using a first order Stochastic Runge Kutta scheme detailed in Ref. (115) with time step *h* = 0.01 chosen to be an order of magnitude smaller than the time to diffuse a Kuhn length, *b*^2^*/*6*D*. We then run the simulation for a Rouse time to allow the polymer to reach steady state. The snapshot in Fig. 2A was taken from a simulation with monomer diffusion coefficient *D*_*n*_ = *D*_0_[1 + *ϵ* cos(2*πn/λ*)], where *λ* = 25 Kuhn lengths and the total chain is 100 Kuhn lengths.

### Self-avoiding, confined polymer simulation details

We also develop more realistic polymer simulations by adapting the polychrom software package, a thin wrapper around OpenMM (116). The deterministic forces applied to the polymer are given by the following potential energy functions detailed in the `polychrom.forces` module: (1) Polymer connectivity via harmonic bonds with energy 0.5*κ*(*r*_*ij*_ − 1)^2^, where *r*_*ij*_ is the distance between the centers of adjacent monomers with diameter *d* = 1, and *κ* is chosen such that the average extension of the bond is 0.1 when the harmonic energy is *k*_*B*_ *T*. (2) Spherical confinement with radius *r*_*C*_ of the form *f*_*C*_ [((*r*_*n*_ − *r*_*C*_)^2^ + *δ*^2^)^1*/*2^ −*δ*] if *r*_*n*_ *> r*_*C*_ and 0 if not, where *r*_*n*_ is the distance of the *n*^th^ monomer to the origin, *f*_*C*_ = 5*k*_*B*_ *T /d* is the confining force, and *δ* = 0.1 is some small number inserted to prevent rounding errors. The confinement energy is thus a smooth version of the function *f*_*C*_ |*r* − *r*_*c*_|, where the radius *r*_*C*_ is chosen such that the total volume fraction of all monomers within the confinement is 0.117. (3) Repulsive short-ranged interactions via a Lennard-Jones like potential:

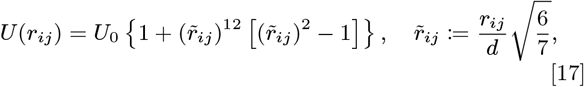

where *U*_0_ = 3*k*_*B*_ *T* represents a finite energy barrier to allow chain passing when *r*_*ij*_ < 0.6*d*. The stochastic forces are implemented according to Eq. (4) using custom Brownian integrators, as detailed in the `contrib/integrators` module of the GitHub repository: https://github.com/open2c/polychrom.

Steady state was determined by running the simulation until the monomer mean squared displacement plateaus at the squared radius of confinement, i.e. until each monomer has had enough time to explore its volume (10^5^ time steps). We then run each simulation for twice this equilibration time and sample 10 steady-state conformations from each run. For each set of parameters, we repeated this procedure over 200 independent simulation runs and thus computed average contact maps from an ensemble of 2000 snapshots. We define a contact as an inter-monomer separation that is less than two monomer diameters.

### Hi-C data processing and compartment identification

To make a meaningful comparison to Hi-C data, we model a region (35-60 Mbp) of Chromosome 2 in murine erythroblasts, a model eukaryotic cell line (80). We then derive the identities of active (A) and inactive (B) monomers in our simulated chain from the data in the following way. First, we iteratively correct the experimental contact map at 100 kilobase resolution such that the rows and columns sum to one, a process that removes experimental biases and ensures equal visibility of all loci (117). We then divide each diagonal of the experimental contact frequency map by its mean in order to produce an “observed over expected” map, which measures structure in the data beyond the average decay of contacts with genomic separation. Subtracting the mean from this “observed over expected” map yields a matrix where positive entries indicate enrichment of contacts above the mean and negative entries denote depletion of contacts below the mean. The first eigenvector (E1) of the resulting map captures the checkerboard pattern characteristic of A/B compartmentalization, and can thus be used to binarize the genome into active (A) and inactive (B) segments. Since the AB identities are determined up to a sign of the entries of E1, we “align” the E1 track to a binned profile of GC content, such that positive entries correlate with active chromatin and negative entries with inactive chromatin (117). Since compartments are typically measured at the 100kb resolution, it is sufficient to assign 4 monomers to each Hi-C bin, i.e. one monomer per 25 kilobases. Note that this resolution is well beyond the persistence length of chromatin, which is on the order of a kilobase (118), justifying the omission of bending rigidity in our simulations.

### Compartment scores

To quantify the degree of compartmentalization observed in both experimental and simulated contact maps, we compute an order parameter—the compartment score. Specifically, we use the definition of the “COMP score 2” introduced in Ref. (84). We first process the simulated contact maps in the same way as the experimental data, i.e. via iterative correction and computation of E1 (see previous section). The rows and columns of the observed over expected map are sorted and binned by quantiles of the E1 track, such that the top left quadrant shows B-B contacts, the bottom right quadrant shows A-A contacts, and the off-diagonal quadrants show contacts between A and B regions. The COMP score is defined by averaging over the top 25% of contacts in each of the 4 quadrants and computing (*AA* + *BB* − 2*AB*)*/*(*AA* + *BB* + 2*AB*). The resulting score is 0.0 if there is no difference between the contact frequencies of same-type and different-type chromatin, and 1.0 if A and B regions are perfectly demixed.

## ACKNOWLEDGMENTS

We thank Richard A. Young, Phillip A. Sharp, Anders S. Hansen, and Leonid Mirny for helpful discussions and for critical reading of the manuscript. This work was supported by the National Science Foundation, through the Biophysics of Nuclear Condensates grant (MCB-2044895) and the Graduate Research Fellowship Program under grant No. 2141064.

## AUTHOR CONTRIBUTIONS

A.G., D.K., M.K., and A.K.C. designed research. A.G. carried out analytical calculations and related numerical implementation. D.K. implemented and carried out Brownian dynamics simulations. All authors discussed and analyzed the results and wrote the manuscript.

## AUTHOR DECLARATION

The authors declare that there are no competing interests. But, for completeness it is noted that A.K.C serves as a consultant (titled Academic Partner) for Flagship Pioneering and its affiliated companies, Apriori Bio and FL72.

## References

1. JN Onuchic, Z Luthey-Schulten, PG Wolynes, Theory of protein folding: The energy land-scape perspective. Annu. Rev. Phys. Chem. 48, 545–600 (1997).

2. VS Pande, AY Grosberg, T Tanaka, Heteropolymer freezing and design: Towards physical models of protein folding. Rev. Mod. Phys. 72, 259–314 (2000).

3. E Shakhnovich, Protein folding thermodynamics and dynamics: Where physics, chemistry, and biology meet. Chem. Rev. 106, 1559–1588 (2006).

4. KA Dill, SB Ozkan, MS Shell, TR Weikl, The protein folding problem. Annu. Rev. Biophys. 37, 289–316 (2008).

5. PG Wolynes, Evolution, energy landscapes and the paradoxes of protein folding. Biochimie 119, 218–230 (2015).

6. R Nassar, GL Dignon, RM Razban, KA Dill, The protein folding problem: The role of theory. J. Mol. Biol. 433, 167126 (2021).

7. JL England, VS Pande, Potential for modulation of the hydrophobic effect inside chaperonins. Biophys. J. 95, 3391–3399 (2008).

8. JL England, Allostery in Protein Domains Reflects a Balance of Steric and Hydrophobic Effects. Structure 19, 967–975 (2011).

9. N Perunov, JL England, Quantitative theory of hydrophobic effect as a driving force of protein structure. Protein Sci. 23, 387–399 (2014).

10. T Misteli, The Self-Organizing Genome: Principles of Genome Architecture and Function. Cell 183, 28–45 (2020).

11. I Solovei, K Thanisch, Y Feodorova, How to rule the nucleus: divide et impera. Curr. Opin. Cell Biol. 40, 47–59 (2016).

12. E Lieberman-Aiden, et al., Comprehensive mapping of long-range interactions reveals folding principles of the human genome. Science 326, 289–293 (2009).

13. M Falk, et al., Heterochromatin drives compartmentalization of inverted and conventional nuclei. Nature 570, 395–399 (2019).

14. B Zhang, PG Wolynes, Topology, structures, and energy landscapes of human chromosomes. Proc. Natl. Acad. Sci. 112, 6062–6067 (2015).

15. B Zhang, PG Wolynes, Shape transitions and chiral symmetry breaking in the energy landscape of the mitotic chromosome. Phys. Rev. Lett. 116, 248101 (2016).

16. M Di Pierro, B Zhang, EL Aiden, PG Wolynes, JN Onuchic, Transferable model for chromo-some architecture. Proc. Natl. Acad. Sci. 113, 12168–12173 (2016).

17. B Zhang, PG Wolynes, Genomic energy landscapes. Biophys. J. 112, 427–433 (2017).

18. G Le Treut, F Képès, H Orland, A polymer model for the quantitative reconstruction of chromosome architecture from hic and gam data. Biophys. J. 115, 2286–2294 (2018).

19. Y Qi, B Zhang, Predicting three-dimensional genome organization with chromatin states. PLOS Comput. Biol. 15, e1007024 (2019).

20. WJ Xie, Y Qi, B Zhang, Characterizing chromatin folding coordinate and landscape with deep learning. PLOS Comput. Biol. 16, e1008262 (2020).

21. JJB Messelink, MCF van Teeseling, J Janssen, M Thanbichler, CP Broedersz, Learning the distribution of single-cell chromosome conformations in bacteria reveals emergent order across genomic scales. Nat. Commun. 12, 1963 (2021).

22. IS Tolokh, NA Kinney, IV Sharakhov, AV Onufriev, Strong interactions between highly-dynamic lamina-associated domains and the nuclear envelope stabilize the 3d architecture of drosophila interphase chromatin. bioRxiv p. 2022.01.28.478236 (2022).

23. GJ Narlikar, R Sundaramoorthy, T Owen-Hughes, Mechanisms and functions of atp-dependent chromatin-remodeling enzymes. Cell 154, 490–503 (2013).

24. F Uhlmann, Smc complexes: from dna to chromosomes. Nat. Rev. Mol. Cell Biol. 17, 399–412 (2016).

25. MA Reid, Z Dai, JW Locasale, The impact of cellular metabolism on chromatin dynamics and epigenetics. Nat. Cell Biol. 19, 1298–1306 (2017).

26. D Zwicker, The intertwined physics of active chemical reactions and phase separation. Curr. Opin. Colloid & Interface Sci. p. 101606 (2022).

27. D Needleman, Z Dogic, Active matter at the interface between materials science and cell biology. Nat. Rev. Mater. 2, 17048 (2017).

28. CP Brangwynne, GH Koenderink, FC MacKintosh, D. Weitz, Intracellular transport by active diffusion. Trends Cell Biol. 19, 423–427 (2009).

29. SC Weber, AJ Spakowitz, JA Theriot, Nonthermal ATP-dependent fluctuations contribute to the in vivo motion of chromosomal loci. Proc. Natl. Acad. Sci. 109, 7338–7343 (2012).

30. A Zidovska, A Weitz David, J Mitchison Timothy, Micron-scale coherence in interphase chromatin dynamics. Proc. Natl. Acad. Sci. 110, 15555–15560 (2013).

31. HA Shaban, R Barth, K Bystricky, Formation of correlated chromatin domains at nanoscale dynamic resolution during transcription. Nucleic Acids Res. 46, e77–e77 (2018).

32. SS Ashwin, T Nozaki, K Maeshima, M Sasai, Organization of fast and slow chromatin revealed by single-nucleosome dynamics. Proc. Natl. Acad. Sci. 116, 19939–19944 (2019).

33. A Javer, et al., Short-time movement of e. coli chromosomal loci depends on coordinate and subcellular localization. Nat. Commun. 4, 3003 (2013).

34. B Gu, et al., Transcription-coupled changes in nuclear mobility of mammalian cis-regulatory elements. Science 359, 1050–1055 (2018).

35. T Germier, et al., Real-Time Imaging of a Single Gene Reveals Transcription-Initiated Local Confinement. Biophys. J. 113, 1383–1394 (2017).

36. R Nagashima, et al., Single nucleosome imaging reveals loose genome chromatin networks via active rna polymerase ii. J. Cell Biol. 218, 1511–1530 (2019).

37. T Nozaki, et al., Dynamic organization of chromatin domains revealed by super-resolution live-cell imaging. Mol. Cell 67, 282–293.e7 (2017).

38. MMC Tortora, H Salari, D Jost, Chromosome dynamics during interphase: a biophysical perspective. Curr. Opin. Genet. & Dev. 61, 37–43 (2020).

39. A Ghosh, NS Gov, Dynamics of active semiflexible polymers. Biophys. J. 107, 1065–1073 (2014).

40. T Eisenstecken, G Gompper, RG Winkler, Conformational properties of active semiflexible polymers. Polymers 8 (2016).

41. T Eisenstecken, A Ghavami, A Mair, G Gompper, RG Winkler, Conformational and dynamical properties of semiflexible polymers in the presence of active noise. AIP Conf. Proc. 1871, 050001 (2017).

42. T Eisenstecken, G Gompper, RG Winkler, Internal dynamics of semiflexible polymers with active noise. The J. Chem. Phys. 146, 154903 (2017).

43. D Osmanović, Y Rabin, Dynamics of active rouse chains. Soft Matter 13, 963–968 (2017).

44. SM Mousavi, G Gompper, RG Winkler, Active brownian ring polymers. The J. Chem. Phys. 150, 064913 (2019).

45. T Saito, T Sakaue, Inferring active noise characteristics from the paired observations of anomalous diffusion. Polymers 11, 2 (2019).

46. SK Anand, SP Singh, Conformation and dynamics of a self-avoiding active flexible polymer. Phys. Rev. E 101, 030501 (2020).

47. A Ghosh, AJ Spakowitz, Statistical behavior of nonequilibrium and living biological systems subjected to active and thermal fluctuations. Phys. Rev. E 105, 014415 (2022).

48. A Ghosh, AJ Spakowitz, Active and thermal fluctuations in multi-scale polymer structure and dynamics. Soft Matter 18, 6629–6637 (2022).

49. RG Winkler, G Gompper, The physics of active polymers and filaments. The J. Chem. Phys. 153, 040901 (2020).

50. N Ganai, S Sengupta, GI Menon, Chromosome positioning from activity-based segregation. Nucleic Acids Res. 42, 4145–4159 (2014).

51. A Awazu, Segregation and phase inversion of strongly and weakly fluctuating brownian particle mixtures and a chain of such particle mixtures in spherical containers. Phys. Rev. E 90, 042308 (2014).

52. SA Sewitz, et al., Heterogeneous chromatin mobility derived from chromatin states is a determinant of genome organisation in <em>s. cerevisiae</em>. bioRxiv p. 106344 (2017).

53. L Liu, G Shi, D Thirumalai, C Hyeon, Chain organization of human interphase chromosome determines the spatiotemporal dynamics of chromatin loci. PLOS Comput. Biol. 14, e1006617 (2018).

54. A Agrawal, N Ganai, S Sengupta, GI Menon, Nonequilibrium biophysical processes influence the large-scale architecture of the cell nucleus. Biophys. J. 118, 2229–2244 (2020).

55. Z Jiang, Y Qi, K Kamat, B Zhang, Phase separation and correlated motions in motorized genome. The J. Phys. Chem. B 126, 5619–5628 (2022).

56. D Loi, S Mossa, LF Cugliandolo, Non-conservative forces and effective temperatures in active polymers. Soft Matter 7, 10193–10209 (2011).

57. AY Grosberg, JF Joanny, Nonequilibrium statistical mechanics of mixtures of particles in contact with different thermostats. Phys. Rev. E 92, 032118 (2015).

58. SN Weber, CA Weber, E Frey, Binary mixtures of particles with different diffusivities demix. Phys. Rev. Lett. 116, 058301 (2016).

59. J Smrek, K Kremer, Small activity differences drive phase separation in active-passive polymer mixtures. Phys. Rev. Lett. 118, 098002 (2017).

60. S Put, T Sakaue, C Vanderzande, Active dynamics and spatially coherent motion in chromo-somes subject to enzymatic force dipoles. Phys. Rev. E 99, 032421 (2019).

61. R Bruinsma, AY Grosberg, Y Rabin, A Zidovska, Chromatin hydrodynamics. Biophys. J. 106, 1871–1881 (2014).

62. I Eshghi, A Zidovska, AY Grosberg, Symmetry-based classification of forces driving chromatin dynamics. Soft Matter 18, 8134–8146 (2022).

63. S Shin, HW Cho, G Shi, D Thirumalai, Transcription-induced active forces suppress chromatin motion by inducing a transient disorder-to-order transition. bioRxiv p. 2022.04.30.490180 (2022).

64. D Saintillan, MJ Shelley, A Zidovska, Extensile motor activity drives coherent motions in a model of interphase chromatin. Proc. Natl. Acad. Sci. 115, 11442–11447 (2018).

65. A Mahajan, W Yan, A Zidovska, D Saintillan, MJ Shelley, Euchromatin activity enhances segregation and compaction of heterochromatin in the cell nucleus. bioRxiv p. 2022.02.22.481494 (2022).

66. V Bianco, E Locatelli, P Malgaretti, Globulelike conformation and enhanced diffusion of active polymers. Phys. Rev. Lett. 121, 217802 (2018).

67. M Foglino, et al., Non-equilibrium effects of molecular motors on polymers. Soft Matter 15, 5995–6005 (2019).

68. I Robles-Rebollo, et al., Cohesin couples transcriptional bursting probabilities of inducible enhancers and promoters. Nat. Commun. 13, 1–16 (2022).

69. E Arner, et al., Transcribed enhancers lead waves of coordinated transcription in transitioning mammalian cells. Science 347, 1010–1014 (2015).

70. J Zhang, et al., Spatial clustering and common regulatory elements correlate with coordinated gene expression. PLOS Comput. Biol. 15, e1006786 (2019).

71. Z Dai, X Dai, Nuclear colocalization of transcription factor target genes strengthens coregulation in yeast. Nucleic Acids Res. 40, 27–36 (2012).

72. V Belcastro, et al., Transcriptional gene network inference from a massive dataset elucidates transcriptome organization and gene function. Nucleic acids research 39, 8677–8688 (2011).

73. M Doi, SF Edwards, The Theory of Polymer Dynamics, International Series of Monographs on Physics. (Clarendon Press, Oxford), (2007).

74. L Harnau, RG Winkler, P Reineker, Dynamic properties of molecular chains with variable stiffness. The J. Chem. Phys. 102, 7750–7757 (1995).

75. O Hallatschek, E Frey, K Kroy, Tension dynamics in semiflexible polymers. Phys. Rev. E 75, 031905 (2007).

76. J Halatek, F Brauns, E Frey, Self-organization principles of intracellular pattern formation. Philos. Transactions Royal Soc. B: Biol. Sci. 373, 20170107 (2018).

77. CA Brackley, et al., Ephemeral protein binding to dna shapes stable nuclear bodies and chromatin domains. Biophys. journal 112, 1085–1093 (2017).

78. TL Tootle, I Rebay, Post-translational modifications influence transcription factor activity: a view from the ets superfamily. Bioessays 27, 285–298 (2005).

79. PE Rouse, A theory of the linear viscoelastic properties of dilute solutions of coiling polymers. The J. Chem. Phys. 21, 1272–1280 (1953).

80. H Zhang, et al., CTCF and transcription influence chromatin structure re-configuration after mitosis. Nat. Commun. 12, 5157 (2021).

81. T Jenuwein, CD Allis, Translating the histone code. Science 293, 1074–1080 (2001).

82. R Margueron, D Reinberg, Chromatin structure and the inheritance of epigenetic information. Nat. Rev. Genet. 11, 285–296 (2010).

83. SL Klemm, Z Shipony, WJ Greenleaf, Chromatin accessibility and the regulatory epigenome. Nat. Rev. Genet. 20, 207–220 (2019).

84. J Nuebler, G Fudenberg, M Imakaev, N Abdennur, LA Mirny, Chromatin organization by an interplay of loop extrusion and compartmental segregation. Proc. Natl. Acad. Sci. 115, E6697–E6706 (2018).

85. MP Backlund, R Joyner, K Weis, WE Moerner, Correlations of three-dimensional motion of chromosomal loci in yeast revealed by the double-helix point spread function microscope. Mol. Biol. Cell 25, 3619 (2014).

86. JJ Parmar, M Woringer, C Zimmer, How the genome folds: The biophysics of four-dimensional chromatin organization. Annu. Rev. Biophys. 48, 231–253 (2019).

87. FS Gnesotto, F Mura, J Gladrow, CP Broedersz, Broken detailed balance and non-equilibrium dynamics in living systems: a review. Reports on Prog. Phys. 81, 066601 (2018).

88. EH Finn, T Misteli, Molecular basis and biological function of variability in spatial genome organization. Science 365, eaaw9498 (2019).

89. T Nagano, et al., Single-cell hi-c reveals cell-to-cell variability in chromosome structure. Nature 502, 59–64 (2013).

90. VI Keizer, et al., Live-cell micromanipulation of a genomic locus reveals interphase chromatin mechanics. Science 377, 489–495 (2022).

91. M Gabriele, et al., Dynamics of ctcf- and cohesin-mediated chromatin looping revealed by live-cell imaging. Science 376, 496–501 (2022).

92. M Socol, et al., Rouse model with transient intramolecular contacts on a timescale of seconds recapitulates folding and fluctuation of yeast chromosomes. Nucleic acids research 47, 6195–6207 (2019).

93. Y Guo, et al., Chromatin jets define the properties of cohesin-driven in vivo loop extrusion. Mol. Cell (2022).

94. CL Wike, et al., Chromatin architecture transitions from zebrafish sperm through early embryogenesis. Genome research 31, 981–994 (2021).

95. NQ Liu, et al., Rapid depletion of ctcf and cohesin proteins reveals dynamic features of chromosome architecture. bioRxiv (2021).

96. BR Parry, et al., The bacterial cytoplasm has glass-like properties and is fluidized by metabolic activity. Cell 156, 183–194 (2014).

97. S Zhang, et al., RNA polymerase II is required for spatial chromatin reorganization following exit from mitosis. Sci. Adv. 7, 1–14 (2021).

98. B van Steensel, EE Furlong, The role of transcription in shaping the spatial organization of the genome. Nat. reviews Mol. cell biology 20, 327–337 (2019).

99. M Di Pierro, DA Potoyan, PG Wolynes, JN Onuchic, Anomalous diffusion, spatial coherence, and viscoelasticity from the energy landscape of human chromosomes. Proc. Natl. Acad. Sci. 115, 7753–7758 (2018).

100. H Salari, M Di Stefano, D Jost, Spatial organization of chromosomes leads to heterogeneous chromatin motion and drives the liquid-or gel-like dynamical behavior of chromatin. Genome Res. 32, 28–43 (2022).

101. VY Goel, MK Huseyin, AS Hansen, Region capture micro-c reveals coalescence of enhancers and promoters into nested microcompartments. bioRxiv (2022).

102. THS Hsieh, et al., Resolving the 3D Landscape of Transcription-Linked Mammalian Chromatin Folding. Mol. Cell 78, 539–553 (2020).

103. A Aljahani, et al., Analysis of sub-kilobase chromatin topology reveals nano-scale regulatory interactions with variable dependence on cohesin and ctcf. Nat. communications 13, 1–13 (2022).

104. S Fanucchi, Y Shibayama, S Burd, M Weinberg, M Mhlanga, Chromosomal contact permits transcription between coregulated genes. Cell 155, 606–620 (2013).

105. TC Voss, GL Hager, Dynamic regulation of transcriptional states by chromatin and transcription factors. Nat. Rev. Genet. 15, 69–81 (2014).

106. A Coulon, CC Chow, RH Singer, DR Larson, Eukaryotic transcriptional dynamics: from single molecules to cell populations. Nat. Rev. Genet. 14, 572–584 (2013).

107. HB Brandão, M Gabriele, AS Hansen, Tracking and interpreting long-range chromatin interactions with super-resolution live-cell imaging. Curr. Opin. Cell Biol. 70, 18–26 (2021).

108. JH Su, P Zheng, SS Kinrot, B Bintu, X Zhuang, Genome-scale imaging of the 3d organization and transcriptional activity of chromatin. Cell 182, 1641–1659 (2020).

109. T Lu, CE Ang, X Zhuang, Spatially resolved epigenomic profiling of single cells in complex tissues. bioRxiv p. 2022.02.17.480825 (2022).

110. AN Boettiger, et al., Super-resolution imaging reveals distinct chromatin folding for different epigenetic states. Nat. 2016 529:7586 529, 418–422 (2016).

111. T Yang, et al., Reconfigurable microbots folded from simple colloidal chains. Proc. Natl. Acad. Sci. 117, 18186–18193 (2020).

112. T Litschel, B Ramm, R Maas, M Heymann, P Schwille, Beating vesicles: Encapsulated protein oscillations cause dynamic membrane deformations. Angewandte Chemie Int. Ed. 57, 16286–16290 (2018).

113. S Christ, T Litschel, P Schwille, R Lipowsky, Active shape oscillations of giant vesicles with cyclic closure and opening of membrane necks. Soft Matter 17, 319–330 (2021).

114. A Goychuk, E Frey, Protein recruitment through indirect mechanochemical interactions. Phys. Rev. Lett. 123, 178101 (2019).

115. AJ Roberts, Modify the improved euler scheme to integrate stochastic differential equations (2012).

116. P Eastman, et al., OpenMM 7: Rapid development of high performance algorithms for molecular dynamics. PLOS Comput. Biol. 13, e1005659 (2017).

117. M Imakaev, et al., Iterative correction of Hi-C data reveals hallmarks of chromosome organization. Nat. Methods 9, 999–1003 (2012).

118. B Beltran, D Kannan, Q MacPherson, AJ Spakowitz, Geometrical heterogeneity dominates thermal fluctuations in facilitating chromatin contacts. Phys. Rev. Lett. 123, 208103 (2019).

