## Supporting Information for "Polymer folding through active processes recreates features of genome organization"

bioRxiv

#### This PDF file includes:

Supporting text
Figs. S1 to S8
SI References

- C Inference of an activity profile that folds the polymer towards a desired conformation 25

### Supporting Information Text

### 1. Theoretical analysis of a generic polymer model

In the following, we recapitulate and generalize the theoretical analysis that was discussed in the main text. In section 1 A "Rouse mode decomposition leads to a non-diagonal system of equations", we first set up the description of a generalized linear polymer that is driven by athermal excitations, and then perform a Rouse mode decomposition. Deriving a framework for the analysis of a generic polymer model will pave the way for the discussion of polymers with specific mechanical properties, see section 2 "Folding patterns predicted by different polymer models". To that end, we derive the steady-state polymer conformation in section 1 B "Derivation of the steady-state polymer conformation". Furthermore, in section 1 C "Coherent motion of the polymer backbone", we relate athermal excitations with a given correlation function to the pairwise correlation between the velocities of different material points along the polymer backbone. In doing so, we draw a possible connection between correlated motion and folding of polymers. Finally, in section 1 D "Conformation of a polymer with a diagonal response matrix", we discuss the conformation of a polymer with translationally invariant mechanical properties. We generalize these results in section 1 E "Perturbation approximation for non-diagonal response matrices" via a perturbation analysis that takes into account weak mechanical inhomogeneities.

**A. Rouse mode decomposition leads to a non-diagonal system of equations.** In the present section, we generalize the formalism presented in the main text to a broader class of continuum polymer models. As discussed in the main text, we idealize the polymer as a space curve $\boldsymbol{r}(s,t)$, where $s \in [-L/2, L/2]$ is a continuous dimensionless material coordinate along the polymer backbone[1]. To drive the polymer dynamics, we explicitly introduce an active random force field $\boldsymbol{\mu}(s,t)$ with zero mean and covariance

$$\langle \boldsymbol{\mu}(s,t) \cdot \boldsymbol{\mu}(s',t') \rangle \coloneqq \mathcal{C}^{\mu}(s,s')\, \delta(t-t')\,, \qquad \text{[S1]}$$

on timescales longer than its decorrelation time. We assume that the solution in which the polymer is embedded is perfectly viscous and has no memory. However, for now we invoke no additional information about the specific mechanical properties of the polymer backbone.

The conservation of momentum implies a balance between all forces that apply at any material point of the polymer,

$$\int ds'\, \mathcal{Q}(s,s') \circ \partial_t \boldsymbol{r}(s',t) = \int ds'\, \mathcal{L}(s,s') \circ \boldsymbol{r}(s',t) + \boldsymbol{\mu}(s,t)\,, \qquad \text{[S2]}$$

where $\mathcal{L}(s,s')$ encodes the elastic properties of the polymer and $\mathcal{Q}(s,s')$ describes viscous couplings. One recovers the dynamics of a Rouse polymer, see section 2 A "Generalized active Rouse polymer", for homogeneous local friction where $\mathcal{Q}(s,s') = \xi\, \delta(s-s')$, and homogeneous line tension where $\mathcal{L}(s,s') = \kappa\, \delta(s-s')\, \partial_{s'}^2$. In general, however, $\mathcal{L}(s,s')$ can also account for pairwise harmonic interactions between different material points, bending rigidity, soft confinement, or material inhomogeneities. Similarly, $\mathcal{Q}(s,s')$ can describe inhomogeneities in drag friction, or even incorporate hydrodynamic coupling between different material points via the Kirkwood-Riseman approximation (1, 2).

[1] The analysis of discrete chains is analogous but involves discrete spectral transforms, which make the calculations longer and less transparent.

We analyze Eq. (S2) with a Rouse mode decomposition (3),

$$\boldsymbol{r}(s,t) \to \tilde{\boldsymbol{r}}_q(t) := \int_{-L/2}^{L/2} ds\, e^{-iqs}\, \boldsymbol{r}(s,t)\,, \tag{S3a}$$

$$\boldsymbol{\mu}(s,t) \to \tilde{\boldsymbol{\mu}}_q(t) := \int_{-L/2}^{L/2} ds\, e^{-iqs}\, \boldsymbol{\mu}(s,t)\,, \tag{S3b}$$

where $L$ is the polymer length. For a compact notation, we concatenate all Rouse modes row-wise into a matrix $\boldsymbol{R}(t)$ with rows $R_{q,\dots}(t) := \tilde{\boldsymbol{r}}_q(t)$. Analogously, we define the random force mode matrix $\boldsymbol{M}(t)$ with rows $M_{q,\dots}(t) := \tilde{\boldsymbol{\mu}}_q(t)$. Finally, we define the active force mode covariance matrix, the friction kernel, and the elastic kernel, respectively[2,3]:

$$\mathcal{C}^\mu(s,s') \to C^\mu_{qk} := \iint ds\, ds'\, e^{-iqs} \mathcal{C}^\mu(s,s') e^{iks'}\,, \tag{S4a}$$

$$\mathcal{Q}(s,s') \to \Xi_{qk} := \iint ds\, ds'\, e^{-iqs} \mathcal{Q}(s,s') e^{iks'}\,, \tag{S4b}$$

$$\mathcal{L}(s,s') \to E_{qk} := \iint ds\, ds'\, e^{-iqs} \mathcal{L}(s,s') e^{iks'}\,. \tag{S4c}$$

Taken together, the dynamics of a polymer with length $L$ are determined by

$$\partial_t \boldsymbol{R}(t) = \Xi^{-1} \cdot \boldsymbol{E} \cdot \boldsymbol{R}(t) + L\, \Xi^{-1} \cdot \boldsymbol{M}(t)\,, \quad \text{where} \quad \left\langle \boldsymbol{M}(t) \cdot \boldsymbol{M}^\dagger(t') \right\rangle = \boldsymbol{C}^\mu\, \delta(t-t')\,. \tag{S5}$$

By identifying the response matrix as $\boldsymbol{J} := -\Xi^{-1} \cdot \boldsymbol{E}$ and the random velocity matrix as $\boldsymbol{H}(t) := L\, \Xi^{-1} \cdot \boldsymbol{M}(t)$, we bring Eq. (S5) into a simpler form that is analogous to the main text,

$$\partial_t \boldsymbol{R}(t) = -\boldsymbol{J} \cdot \boldsymbol{R}(t) + \boldsymbol{H}(t)\,, \tag{S6}$$

where the response matrix $\boldsymbol{J}$ must be positive definite for the system to remain stable. As discussed in the main text and in section 2 A.3 "Activity modulations lead to bending", the response matrix is diagonal for a Rouse polymer, $J_{qk} = \xi^{-1}\kappa\, q^2\, \delta_{qk}$, and in general for any polymer with homogeneous mechanical properties. In the limit of infinitely long polymers ($L \to \infty$), one has $L\delta_{qk} \to 2\pi\, \delta(q-k)$ and the inner product becomes $L^{-1}\,[\boldsymbol{J} \cdot \boldsymbol{R}(t)]_{qk} := \int \frac{dp}{2\pi}\, J_{qp} \cdot R_{pk}(t)$. The covariance of the random velocity matrix,

$$\left\langle \boldsymbol{H}(t) \cdot \boldsymbol{H}^\dagger(t') \right\rangle := \boldsymbol{C}\, \delta(t-t')\,, \quad \text{with} \quad \boldsymbol{C} := L^2\, \Xi^{-1} \cdot \boldsymbol{C}^\mu \cdot \Xi^{-1,\dagger}\,, \tag{S7}$$

illustrates, as discussed in section 1 C "Coherent motion of the polymer backbone", how correlated motion can arise from correlated active forces or viscous couplings.

***A.1. Response matrix of a discrete Rouse polymer depends on chain topology.*** For completeness, we now discuss the mechanical properties of a discrete chain with constant spring stiffness (tension) and friction coefficient. In contrast to an infinitely long continuous polymer where the Rouse mode decomposition amounts to a continuous Fourier transform, for a discrete chain we need to take into account the topology of the polymer and the corresponding boundary conditions. Specifically, a closed ring-shaped Rouse polymer of length $N$ is decomposed by a discrete Fourier transform with

[2] These matrices become finer grained as smaller nontrivial wave modes enter with increasing polymer length $L$, and turn into continuous fields for $L \to \infty$.

[3] While these matrices are infinitely large for continuous polymers, they have a finite size for discrete polymers. The matrix size then encodes the small wavelength cutoff (largest admissible wave mode).

discrete wave coefficients $n \in [0, N-1]$, while an open Rouse polymer requires a discrete cosine transform. Moreover, these two polymer topologies lead to different response matrices,

$$J_{nm} = \frac{\kappa}{\xi}\,\delta_{nm} \times \begin{cases} \left[2\sin\left(\frac{\pi n}{N}\right)\right]^2, & \text{for a closed ring, or} \\ \left[2\sin\left(\frac{\pi n}{2N}\right)\right]^2, & \text{for an open polymer.} \end{cases} \qquad \text{[S8]}$$

Overall, especially in the context of model extensions, it is much simpler to study an infinitely long continuous polymer than to study a discrete chain.

**B. Derivation of the steady-state polymer conformation.** To analyze the conformation of the polymer, we turn to the moments of the Rouse matrix. The first moment of the Rouse matrix vanishes for zero-mean excitations, $\langle \boldsymbol{R}(t)\rangle = 0$, cf. Eq. (S6). For Gaussian excitations, the polymer conformation is therefore entirely captured by the second moment of the Rouse matrix, $\langle \boldsymbol{R}(t)\cdot\boldsymbol{R}^\dagger(t')\rangle$. To find a constitutive equation for the second moment of the Rouse matrix, we multiply Eq. (S6) with $\boldsymbol{R}^\dagger(t')$ from the right and average over different trajectories:

$$\partial_t \left\langle \boldsymbol{R}(t)\cdot\boldsymbol{R}^\dagger(t')\right\rangle = -\boldsymbol{J}\cdot\left\langle \boldsymbol{R}(t)\cdot\boldsymbol{R}^\dagger(t')\right\rangle + \left\langle \boldsymbol{H}(t)\cdot\boldsymbol{R}^\dagger(t')\right\rangle\,. \qquad \text{[S9]}$$

To proceed, we need to determine the *driving term* $\langle \boldsymbol{H}(t)\cdot\boldsymbol{R}^\dagger(t')\rangle$ of this non-homogeneous matrix differential equation (S9). To that end, we formally integrate the conjugate transpose of Eq. (S6) forward in time,

$$\boldsymbol{R}^\dagger(t') = \int_{-\infty}^{t'} dt'' \left[-\boldsymbol{R}^\dagger(t'')\cdot\boldsymbol{J}^\dagger + \boldsymbol{H}^\dagger(t'')\right]. \qquad \text{[S10]}$$

After multiplying Eq. (S10) with the Rouse matrix $\boldsymbol{H}(t)$ from the left, averaging over different trajectories, and substituting the covariance of the random velocity matrix, Eq. (S7), one then has:

$$\left\langle \boldsymbol{H}(t)\cdot\boldsymbol{R}^\dagger(t')\right\rangle = -\int_{-\infty}^{t'} dt'' \left\langle \boldsymbol{H}(t)\cdot\boldsymbol{R}^\dagger(t'')\right\rangle\cdot\boldsymbol{J}^\dagger + \boldsymbol{C}\,\Theta(t'-t)\,, \qquad \text{[S11]}$$

where we have defined the Heaviside step function $\Theta(x) := 1\,\forall\,x > 0$, else zero. The matrix integral equation (S11) is solved by

$$\left\langle \boldsymbol{H}(t)\cdot\boldsymbol{R}^\dagger(t')\right\rangle = \Theta(t'-t)\,\boldsymbol{C}\cdot e^{-\boldsymbol{J}^\dagger\,(t'-t)}\,, \qquad \text{[S12]}$$

which we substitute into the constitutive equation (S9) for the second moment of the Rouse matrix. Note that Eq. (S12) vanishes for all times after a given reference time, $t \geq t'$, which means that excitations are independent of the past polymer conformation (causality). Therefore, it follows from Eq. (S9) that the second Rouse moment is given by

$$\left\langle \boldsymbol{R}(t)\cdot\boldsymbol{R}^\dagger(t')\right\rangle = e^{-\boldsymbol{J}\,(t-t')}\cdot\boldsymbol{X}(t')\,,\quad \text{for}\quad t \geq t'\,, \qquad \text{[S13]}$$

where $\boldsymbol{X}(t)$ is a Hermitian matrix. We also find an expression for the second Rouse moment for all times that precede a given reference time, $t \leq t'$, by taking the conjugate transpose of Eq. (S13) and exchanging the time labels $t \leftrightarrow t'$:

$$\left\langle \boldsymbol{R}(t)\cdot\boldsymbol{R}^\dagger(t')\right\rangle = \boldsymbol{X}(t)\cdot e^{-\boldsymbol{J}^\dagger\,(t'-t)}\,,\quad \text{for}\quad t \leq t'\,. \qquad \text{[S14]}$$

To now determine the Hermitian matrix $\boldsymbol{X}(t)$, we insert the second moment of the Rouse matrix, Eq. (S14), back into the constitutive equation (S9) for the second moment of the Rouse matrix, while studying *past* times $t \leq t'$. In doing so, we obtain the Lyapunov equation as consistency relation,

$$\partial_t \boldsymbol{X}(t) + \boldsymbol{X}(t) \cdot \boldsymbol{J}^\dagger + \boldsymbol{J} \cdot \boldsymbol{X}(t) = \boldsymbol{C}\,, \qquad \text{[S15]}$$

which is well known from control theory (4) and has the following steady-state solution, i.e. $\partial_t \boldsymbol{X}(t) = 0$, in the long time limit (5):

$$\lim_{t\to\infty} \boldsymbol{X}(t) = \int_0^\infty d\tau\, e^{-\boldsymbol{J}\tau} \cdot \boldsymbol{C} \cdot e^{-\boldsymbol{J}^\dagger \tau}\,. \qquad \text{[S16]}$$

Henceforth, we indicate the steady-state polymer conformation with the abbreviation $\boldsymbol{X} \equiv \lim_{t\to\infty} \boldsymbol{X}(t)$. Finally transforming back into real space allows us to determine the mean squared separation and the tangent-tangent autocorrelation between pairs of material points. Our closed analytical expressions illustrate how athermal excitations that break translational invariance induce an effective coupling between different mechanical modes, which can give rise to *patterns* in the polymer conformation.

**C. Coherent motion of the polymer backbone.** In this section, we seek the velocity correlation function $\mathcal{C}^v(s,s') = \langle \boldsymbol{v}(s,t) \cdot \boldsymbol{v}(s',t)\rangle$, where $\boldsymbol{v}(s,t) := \partial_t \boldsymbol{r}(s,t)$ is the realized velocity of a given material point. To compute this velocity, we consider the finite difference approximation with a finite measurement time window $\Delta t$. In Fourier space, the velocity correlation is then given by

$$\boldsymbol{C}^v := \left\langle \frac{\boldsymbol{R}(t+\Delta t) - \boldsymbol{R}(t)}{\Delta t} \cdot \frac{\boldsymbol{R}^\dagger(t+\Delta t) - \boldsymbol{R}^\dagger(t)}{\Delta t} \right\rangle. \qquad \text{[S17]}$$

By substituting the second moment of the Rouse matrix, Eq. (S13) and Eq. (S14), we obtain:

$$\boldsymbol{C}^v = \frac{1}{\Delta t^2}\left[2\boldsymbol{X} - e^{-\boldsymbol{J}\Delta t} \cdot \boldsymbol{X} - \boldsymbol{X} \cdot e^{-\boldsymbol{J}^\dagger \Delta t}\right] \approx \frac{1}{\Delta t}\left[\boldsymbol{J} \cdot \boldsymbol{X} + \boldsymbol{X} \cdot \boldsymbol{J}^\dagger\right]. \qquad \text{[S18]}$$

At long times where the polymer conformation has reached a steady state, $\partial_t \boldsymbol{X}(t) = 0$, a comparison with the Lyapunov equation (S15) yields:

$$\boldsymbol{C}^v = \frac{\boldsymbol{C}}{\Delta t}\,, \quad \text{which in sequence space implies} \quad \mathcal{C}^v(s,s') = \frac{\mathcal{C}(s,s')}{\Delta t}\,. \qquad \text{[S19]}$$

Thus, spatial correlations between the athermal excitations at different material points will lead to correlated motion of the polymer backbone.

**D. Conformation of a polymer with a diagonal response matrix.** Polymers with homogeneous material properties, such as line tension and friction, have a diagonal response matrix $J_{qk} \equiv J_{qq}\,\delta_{qk}$. For such a diagonal response matrix, the correlation between Rouse modes is given by [cf. Eq. (S16)]:

$$X_{qk} = \frac{C_{qk}}{J_{qq} + J_{kk}}\,. \qquad \text{[S20]}$$

In general, however, the response matrix will be non-diagonal and we need to use a perturbation approach to approximate the matrix exponentials in Eq. (S16).

**E. Perturbation approximation for non-diagonal response matrices.** To evaluate Eq. (S16) for non-diagonal response matrices, we make use of several approximations. First and foremost, we require that the material properties of the polymer are dominated by topological connectivity between adjacent material points and by homogeneous mechanical features of the polymer backbone. These translationally invariant properties are encoded in the diagonal entries of the response matrix,

$$\bar{J}_{qk} := \begin{cases} J_{qk}, & q = k\,, \\ 0, & q \neq k\,, \end{cases} \tag{S21}$$

which we refer to as *dominant homogeneous contribution.* In addition, we permit weak mechanical inhomogeneities that lead to off-diagonal entries in the response matrix:

$$\delta J_{qk} := \begin{cases} 0, & q = k\,, \\ J_{qk}, & q \neq k\,. \end{cases} \tag{S22}$$

By establishing this hierarchy of dominant diagonal entries and weak off-diagonal entries[4] , $\delta J_{qk} \ll \bar{J}_{qk}$, the response matrix

$$\boldsymbol{J} = \bar{\boldsymbol{J}} \cdot \left[\boldsymbol{I} + \bar{\boldsymbol{J}}^{-1} \cdot \delta \boldsymbol{J}\right], \quad \text{reveals} \quad \left[\bar{\boldsymbol{J}}^{-1} \cdot \delta \boldsymbol{J}\right]_{qk} \ll 1\,, \tag{S23}$$

as expansion parameter that we can use for approximations ; $\boldsymbol{I}$ is the identity matrix. To approximate the matrix exponentials in Eq. (S16), we use a perturbation formula (6),

$$e^{-\boldsymbol{J}t} \approx e^{-\bar{\boldsymbol{J}}t} \cdot \left[\boldsymbol{I} - \boldsymbol{\Lambda}(t)\right], \tag{S24}$$

where the first-order correction is given by

$$\Lambda_{qk}(t) = \delta J_{qk}\, \frac{e^{(J_{qq}-J_{kk})t} - 1}{J_{qq} - J_{kk}}\,. \tag{S25}$$

For simplicity, in the present work we do not consider higher order corrections. Next, we decompose the matrix of correlations between different excitation modes into (i) statistically independent excitations with homogeneous magnitude $C_0$, and (ii) statistically correlated excitations:

$$\boldsymbol{C} = C_0 \left[\boldsymbol{I} + \delta\widehat{\boldsymbol{C}}\right], \tag{S26}$$

where $\delta\widehat{\boldsymbol{C}}$ is a dimensionless matrix. Substituting Eq. (S26) and Eq. (S24) into Eq. (S16), gives to lowest order:

$$\boldsymbol{X} \approx C_0 \int_0^\infty dt\, e^{-\bar{\boldsymbol{J}}t} \cdot \left[\boldsymbol{I} + \delta\widehat{\boldsymbol{C}} - \boldsymbol{\Lambda}(t) - \boldsymbol{\Lambda}^\dagger(t)\right] \cdot e^{-\bar{\boldsymbol{J}}t}\,, \tag{S27}$$

where the first two terms in the squared brackets reproduce Eq. (S20) in the limit $\boldsymbol{\Lambda} \to 0$. Finally, substituting Eq. (S25) into Eq. (S27), and carrying out the integration, results in

$$X_{qk} = \frac{C_{qk}}{J_{qq} + J_{kk}} - \frac{1}{2}\, \frac{C_0}{J_{qq} + J_{kk}} \left[\frac{\delta J_{qk}}{J_{kk}} + \frac{\delta J^*_{kq}}{J_{qq}}\right]. \tag{S28}$$

We use this linear expansion in section 2 A.1 "Friction modulations do not affect conformation" to show that modulations in the friction coefficient of different material points do not affect the polymer conformation. Furthermore, in section 2 A.2 "Tension modulations do not induce folding", we also show that modulations in line tension do not lead to polymer folding.

[4] Note that, in the Methods section of the main text and in section 2 B "Passive Rouse polymer with weak long-ranged harmonic interactions", we make identical arguments to derive a a matrix of weak long-ranged harmonic interactions that can fold a passive polymer into specific conformations.

**F. Mean squared distance traveled by a specific polymer locus.** An important dynamic observable, in addition to the pairwise velocity correlation function that we have studied in section 1 C "Coherent motion of the polymer backbone", is the mean squared distance

$$\left\langle [\boldsymbol{r}(s,t) - \boldsymbol{r}(s,t')]^2 \right\rangle = \langle \boldsymbol{r}(s,t) \cdot \boldsymbol{r}(s,t) \rangle + \langle \boldsymbol{r}(s,t') \cdot \boldsymbol{r}(s,t') \rangle - 2 \langle \boldsymbol{r}(s,t) \cdot \boldsymbol{r}(s,t') \rangle \ , \qquad \text{[S29]}$$

which a given polymer locus travels within some time window $t - t'$. We transform the correlation between Rouse modes at different times, Eq. (S13), back into real space and use the same perturbation approximation as outlined above, to find

$$\left\langle [\boldsymbol{r}(s,t+\Delta t) - \boldsymbol{r}(s,t)]^2 \right\rangle = 2 \sum_{qk} e^{i(q-k)s} \left[ \left(1 - e^{-J_{qq}\Delta t}\right) X_{qk} + \sum_n e^{-J_{qq}\Delta t} \Lambda_{qn}(\Delta t) X_{nk} \right] . \qquad \text{[S30]}$$

Next, we substitute the definition Eq. (S25) of the linear correction term $\Lambda_{qn}(\Delta t) \propto \delta J_{qn}$. Moreover, we also substitute the steady-state correlation between Rouse modes, Eq. (S28), only keep terms up to linear order in $\delta \widehat{C}_{qk}$ or $\delta J_{qk}$, and assume a Hermitian response matrix:

$$\begin{aligned} \left\langle [\boldsymbol{r}(s,t+\Delta t) - \boldsymbol{r}(s,t)]^2 \right\rangle = \sum_{qk} e^{i(q-k)s} \Bigg\{ & \left(1 - e^{-J_{qq}\Delta t}\right) \frac{C_0 \, \delta_{qk}}{J_{qq}} + \left(1 - e^{-J_{qq}\Delta t}\right) \frac{2 C_0 \, \delta \widehat{C}_{qk}}{J_{qq} + J_{kk}} \\ & - \frac{C_0 \, \delta J_{qk}}{J_{qq} J_{kk}} \left[ 1 - \frac{J_{qq} e^{-J_{kk}\Delta t} - J_{kk} e^{-J_{qq}\Delta t}}{J_{qq} - J_{kk}} \right] \Bigg\} + \mathcal{O}(\delta J^2) + \mathcal{O}(\delta J \, \delta \widehat{C}) \, . \qquad \text{[S31]} \end{aligned}$$

In the following sections, we use this result to quantify the mean squared distance that a specific polymer locus in an active or a passive polymer travels within some time window.

### 2. Folding patterns predicted by different polymer models

In the following, we discuss the two specific polymer models that we introduced in the main text. First, in section 2 A "Generalized active Rouse polymer", we recapitulate the Rouse model that we used in the main text and generalize it by permitting modulations in line tension and in friction coefficient. Then, in section 2 B "Passive Rouse polymer with weak long-ranged harmonic interactions", we discuss a Rouse model where polymer folding is induced by weak long-ranged harmonic interactions.

**A. Generalized active Rouse polymer.** Adjacent material points of the polymer interact through Hookean springs, which can in principle vary in stiffness $\kappa(s)$. These harmonic interactions give rise to line tension and a Laplace pressure proportional to the local curvature. When the polymer moves through the surrounding fluid, each curve segment experiences a drag friction $\xi(s)$ that can also vary along the polymer backbone. We neglect hydrodynamic coupling between different material points, which can be introduced in future studies through the Kirkwood-Riseman approximation (1, 2). Taking these deterministic mechanisms together, the Langevin dynamics of the polymer are given by

$$\partial_t \boldsymbol{r}(s,t) = \Big[\xi(s)\Big]^{-1} \partial_s \Big[\kappa(s)\, \partial_s \boldsymbol{r}(s,t)\Big] + \boldsymbol{\eta}(s,t)\,, \tag{S32}$$

where $\boldsymbol{\eta}(s,t)$ is a Gaussian random displacement velocity field with zero mean. Analogously to the main text, the athermal excitations $\boldsymbol{\eta}(s,t)$ are characterized by the following correlation function:

$$\langle \boldsymbol{\eta}(s,t) \cdot \boldsymbol{\eta}(s',t') \rangle \coloneqq \mathcal{C}(s,s')\, \delta(t-t')\,. \tag{S33}$$

Within the formalism introduced in section 1 A "Rouse mode decomposition leads to a non-diagonal system of equations", the friction operator is given by $\mathcal{Q}(s,s') = \xi(s)\, \delta(s-s')$ and the elastic operator is given by $\mathcal{L}(s,s') = \delta(s-s')\, \partial_{s'}[\kappa(s')\partial_{s'}]$. In the following, we analyze several limiting cases of this model.

***A.1. Friction modulations do not affect conformation.*** We first keep the spring stiffness (tension) fixed, $\kappa(s) = \bar{\kappa}$, and consider the effect of small friction modulations $\xi(s) = \bar{\xi}\,[1+\epsilon(s)]$ around an average value of $\bar{\xi}$, with $\epsilon(s) \ll 1$. The spectrum of these friction modulations is determined by the Fourier transform $\epsilon(s) \to \epsilon_q$, as defined by Eq. (S3). Then, the response matrix is given by $J_{qk} \approx \bar{\xi}^{-1}\, \bar{\kappa}\, [q^2\, \delta_{qk} - L^{-1}\, k^2\, \epsilon_{q-k}]$ with $\epsilon_0 = 0$. Furthermore, for homogeneous activity $A_0$ one has $\mathcal{C}(s,s') = [\xi(s)]^{-1}\, A_0\, \delta(s-s')$, which in Fourier space corresponds to $C_{qk} \approx \bar{\xi}^{-1}\, A_0\, [L\delta_{qk} - \epsilon_{q-k}]$. Taken together, we finally evaluate Eq. (S28) to determine the steady-state polymer conformation:

$$X_{qk} = b^2\, \frac{L\delta_{qk}}{q^2}\,, \tag{S34}$$

where we have defined the characteristic length $b \coloneqq \sqrt{A_0/(2\bar{\kappa})}$ in analogy to the main text. Taking the limit of an infinitely long polymer, $L \to \infty$ so that $L\delta_{qk} \to 2\pi\, \delta(q-k)$, and transforming Eq. (S34) back into real space, we find that the steady state conformation of the polymer is characterized by

$$\langle \boldsymbol{r}(s,t) \cdot \boldsymbol{r}(s',t) \rangle = -\frac{1}{2}\, b^2\, \|s-s'\|\,, \tag{S35}$$

which is identical to a Rouse polymer with homogeneous friction coefficient. This result shows that, to linear order, friction modulations do not change the conformation of a Rouse polymer [Fig. S1A,E].

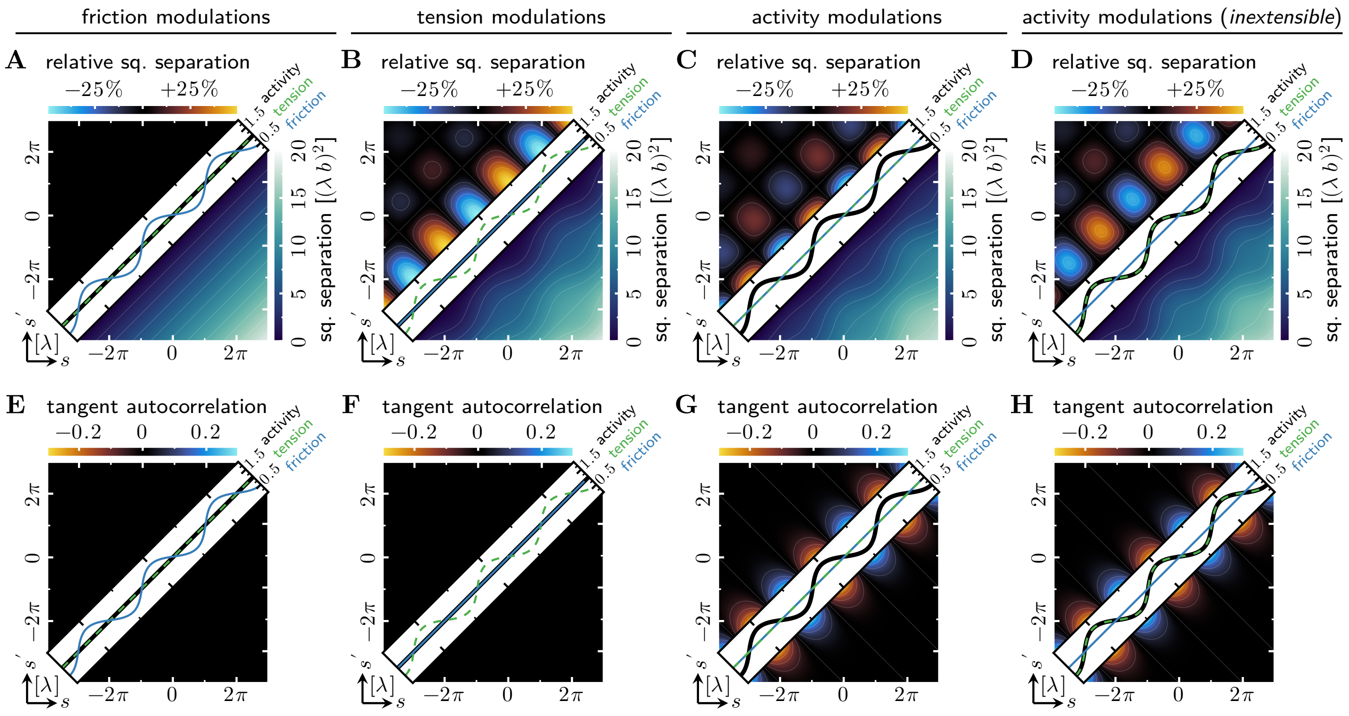

Fig. S1. **Effect of different types of inhomogeneities (diagonal panels) on the conformation of a Rouse polymer. A-D)** *Bottom right panels:* Absolute mean squared separation between different material points. *Top left panels:* Relative mean squared separation when compared to a homogeneous Rouse polymer. **E-H)** The correlation between the tangent vectors at different material points is an indicator of polymer shape. **A,E)** A Rouse polymer that is subject to friction modulations has the same conformation as a homogeneous Rouse polymer. **B,F)** A Rouse polymer that is subject to tension modulations shows variations in the mean squared separation between different material points, but has the same shape as a homogeneous Rouse polymer. **C,G)** A Rouse polymer that is subject to activity modulations changes its conformation and shape. Panels are identical to Fig. 2B,C in the main text. **D,H)** A Rouse polymer that is subject to activity modulations and tension modulations to enforce inextensibility (on average), has the same shape as the corresponding extensible Rouse polymer (compare with panel *G*). The relative squared separation shows that the *boundaries* of active segments come closer together, while the boundaries of inactive segments move farther apart, as one would expect from induced bending. Furthermore, the relative squared separation also shows that, as one would expect from an inextensible polymer, local ($s \sim s'$) stretching and contraction are suppressed.

***A.2. Tension modulations do not induce folding.*** Next, we keep the friction coefficient of each material point fixed, $\xi(s) = \bar{\xi}$, and consider the effect of small spring stiffness (tension) modulations $\kappa(s) = \bar{\kappa}\,[1 + \epsilon(s)]$ around an average value of $\bar{\kappa}$, with $\epsilon(s) \ll 1$. The spectrum of these tension modulations is determined by the Fourier transform $\epsilon(s) \to \epsilon_q$, as defined by Eq. (S3). Then, the response matrix is given by $J_{qk} = \bar{\xi}^{-1}\,\bar{\kappa}\,[q^2\,\delta_{qk} + L^{-1}\,qk\,\epsilon_{q-k}]$ with $\epsilon_0 = 0$. Furthermore, for homogeneous activity one has $\mathcal{C}(s,s') = \bar{\xi}^{-1}\,A_0\,\delta(s-s')$, which in Fourier space corresponds to $C_{qk} = \bar{\xi}^{-1}\,A_0\,L\delta_{qk}$. Taken together, we finally evaluate Eq. (S28) to determine the steady-state polymer conformation:

$$X_{qk} = b^2 \left[ \frac{L\delta_{qk}}{q^2} - \frac{\epsilon_{q-k}}{qk} \right] , \qquad \text{[S36]}$$

where the characteristic length is given by $b := \sqrt{A_0/(2\bar{\kappa})}$, as before. In the limit of an infinitely long polymer, $L \to \infty$ so that $L\delta_{qk} \to 2\pi\,\delta(q-k)$, the correlation between tangent vectors $\boldsymbol{\tau}(s,t) := b^{-1}\,\partial_s \boldsymbol{r}(s,t)$ at different material points is given by

$$\langle \boldsymbol{\tau}(s,t) \cdot \boldsymbol{\tau}(s',t) \rangle = \Big[1 - \epsilon(s)\Big]\,\delta(s-s')\,. \qquad \text{[S37]}$$

Note that Eq. (S37) reduces to $\langle \boldsymbol{\tau}(s,t) \cdot \boldsymbol{\tau}(s',t) \rangle = \delta(s-s')$ if one defines a position-dependent characteristic length $b(s) := \sqrt{A_0/[2\kappa(s)]}$. This result shows that tension modulations alone do not introduce long-ranged correlations and thus cannot fold the polymer [Fig. S1F]. Instead, tension modulations only lead to local stretching and contraction of the polymer backbone.

Transforming Eq. (S36) back into real space quantifies the effect of local stretching and contraction on the correlation between the positions of different material points:

$$\langle \boldsymbol{r}(s,t) \cdot \boldsymbol{r}(s',t) \rangle = -\frac{1}{2}\, b^2 \left[ \| s - s' \| + \int dx\, \epsilon(x)\, G^{\kappa}(s-x, s'-x) \right] , \qquad \text{[S38]}$$

with the following Green's function kernel:

$$G^{\kappa}(s_1, s_2) = \frac{1}{2}\, \mathrm{sgn}(s_1)\, \mathrm{sgn}(s_2)\, . \qquad \text{[S39]}$$

Here, $\mathrm{sgn}(x)$ refers to the sign function. The local stretching and contraction of the polymer backbone, cf. Eq. (S37), results in modulations on the mean squared separation between different material points [Fig. S1B].

***A.3. Activity modulations lead to bending.*** So far, we have seen that neither tension modulations nor friction modulations can fold a Rouse polymer. Now, we keep the friction coefficient of each material point fixed, $\xi(s) = \bar{\xi}$, and also consider homogeneous line tension, $\kappa(s) = \bar{\kappa}$. Then, the response matrix is given by $J_{qk} = \bar{\xi}^{-1}\, \bar{\kappa}\, q^2\, \delta_{qk}$, and is thus diagonal. Instead of introducing mechanical inhomogeneities, we turn towards modulations in the magnitude of statistically independent athermal excitations, $\mathcal{C}(s,s') = \bar{\xi}^{-1}\, A_0\, [1 + \epsilon(s)]\, \delta(s-s')$, in the following referred to as "activity modulations". The spectrum of these activity modulations is determined by the Fourier transform $\epsilon(s) \to \epsilon_q$ as defined by Eq. (S3), so that one has $C_{qk} = \bar{\xi}^{-1}\, A_0\, [L\delta_{qk} + \epsilon_{q-k}]$ where $\epsilon_0 = 0$. Taken together, we finally evaluate Eq. (S20) to determine the steady-state polymer conformation:

$$X_{qk} = b^2 \left[ \frac{L\delta_{qk}}{q^2} + \frac{2\,\epsilon_{q-k}}{q^2 + k^2} \right] , \qquad \text{[S40]}$$

where the characteristic length is given by $b := \sqrt{A_0/(2\bar{\kappa})}$, as before.

Taking the limit of an infinitely long polymer, $L \to \infty$ so that $L\delta_{qk} \to 2\pi\, \delta(q-k)$, and transforming Eq. (S40) back into real space quantifies the effect of activity modulations on the conformation of a Rouse polymer:

$$\langle \boldsymbol{r}(s,t) \cdot \boldsymbol{r}(s',t) \rangle = -\frac{1}{2}\, b^2 \left[ \| s - s' \| + \int dx\, \epsilon(x)\, G^{A}(s-x, s'-x)\, , \right] \qquad \text{[S41]}$$

with the following Green's function kernel:

$$G^{A}(s_1, s_2) = \frac{1}{\pi} \left[ 2\gamma_e + \log\left( s_1^2 + s_2^2 \right) \right] . \qquad \text{[S42]}$$

Here, $\gamma_e$ refers to the Euler–Mascheroni constant. We use these results to plot the mean squared separation between different material points for a Rouse polymer driven by activity modulations with a sinusoidal profile [Fig. S1C] and with a step profile [Fig. S2A].

One can now use $\partial_s \partial_{s'} G^A(s,s')$ to calculate the correlation between tangent vectors $\boldsymbol{\tau}(s,t) := b^{-1}\, \partial_s \boldsymbol{r}(s,t)$ at different material points. However, in the present context we can learn more from

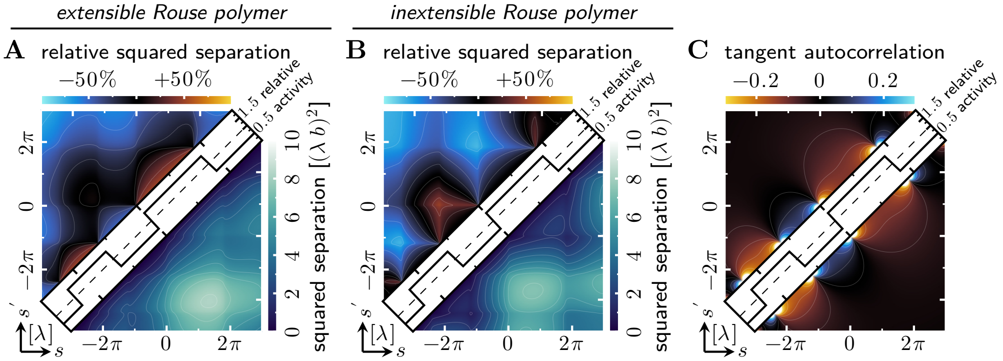

Fig. S2. **Sharp modulations in activity lead to box-like features in the mean squared separation map. A)** *Bottom right panel:* Absolute mean squared separation between different material points of an *extensible* Rouse polymer driven by statistically independent excitations with inhomogeneous magnitude. Note that the mean squared separation can change non-monotonically with distance in sequence space, $s - s'$. *Top left panel:* Relative mean squared separation when compared to a homogeneous Rouse polymer. **B)** *Bottom right panel:* Absolute mean squared separation between different material points of an *inextensible* Rouse polymer driven by statistically independent excitations with inhomogeneous magnitude. *Top left panel:* Relative mean squared separation when compared to a homogeneous polymer. The relative mean squared separation shows features of bending (boundaries of more active polymer segments come closer together) and straightening (boundaries of less active polymer segments move farther apart). **C)** The extensible and the inextensible Rouse polymer have the same steady-state shape, as characterized by the correlation between the tangent vectors at different material points. Polymer segments with an increased activity exhibit bending (anti-alignment of the tangent vectors) while polymer segments with a decreased activity show straightening (alignment of the tangent vectors).

a semi-spectral representation that we obtain by transforming $qkX_{qk}$ back into real space [cf. Eq. (S40)]:

$$\langle \boldsymbol{\tau}(s,t) \cdot \boldsymbol{\tau}(s',t) \rangle = [1 + \epsilon(s)]\, \delta(s - s') - \int \frac{dq}{2\pi} \left[ \epsilon_q \, \frac{\|q\|}{2} \, e^{-\frac{\|q\|}{2}\|s-s'\|} \right] e^{iq\frac{s+s'}{2}} \, . \qquad \text{[S43]}$$

The first term in Eq. (S43) shows that modulations in activity change the local contour length of the polymer backbone, thereby leading to stretching and contraction. A similar effect also arises from modulations in tension, cf. Eq. (S37), and is therefore not a characteristic feature of polymer bending. Instead, one can understand the local activity-induced stretching and contraction by considering independent Rouse polymer segments that are sufficiently short, so that the activity profile within each segment is almost homogeneous. In contrast to a Rouse polymer with homogeneous activity, however, we find that different polymer segments are *not* independent, as the second term in Eq. (S43) shows. Specifically, modulations in activity induce an effective coupling between distinct segments of the Rouse polymer, thus changing its conformation through bending [Figs. S1G and S2C].

***A.4. Enforcing inextensibility in active polymers.*** In the previous sections, we have seen that modulations in activity and in spring stiffness (line tension) along the backbone of a Rouse polymer both induce local changes in the contour length of each polymer segment. To disentangle changes in shape from changes in contour length, we will now enforce inextensibility of the polymer backbone. To that end, we use the line tension as a Lagrange multiplier field $\kappa(s)$ that enforces a local conservation law for the length of each segment (7).

In an active Rouse polymer that is subject to statistically independent athermal excitations with modulations in magnitude, $\mathcal{C}(s,s') = \bar{\xi}^{-1} A_0 \, [1 + \epsilon(s)]\, \delta(s - s')$, the polymer backbone undergoes stretching and contraction as quantified by the first term in Eq. (S43). For these strain modulations to be balanced out by local line tension, cf. Eq. (S37), this line tension must be proportional to the local level of activity, $\kappa(s) = \bar{\kappa}\,[1 + \epsilon(s)]$. Then, the correlation between tangent vectors $\boldsymbol{\tau}(s,t) \coloneqq b^{-1}\,\partial_s \boldsymbol{r}(s,t)$ at different material points is given by

$$\langle \boldsymbol{\tau}(s,t) \cdot \boldsymbol{\tau}(s',t) \rangle = \delta(s - s') - \int \frac{dq}{2\pi} \left[ \epsilon_q \, \frac{\|q\|}{2} \, e^{-\frac{\|q\|}{2}\,\|s-s'\|} \right] e^{iq\frac{s+s'}{2}} \,, \qquad \text{[S44]}$$

where the first term on the right-hand side indicates the local conservation of length. Taken together, we finally evaluate Eq. (S28) to determine the steady-state conformation of the polymer:

$$X_{qk} = b^2 \left[ \frac{L\delta_{qk}}{q^2} + \epsilon_{q-k} \left( \frac{2}{q^2 + k^2} - \frac{1}{qk} \right) \right] , \qquad \text{[S45]}$$

where the characteristic length is given by $b \coloneqq \sqrt{A_0/(2\bar{\kappa})}$, as before. Using Eq. (S45) and taking the limit of an infinitely long polymer, $L \to \infty$ so that $L\delta_{qk} \to 2\pi\,\delta(q-k)$, we thus determine the steady-state conformation of an inextensible active Rouse polymer. As expected, local stretching and contraction of the polymer backbone are suppressed by enforcing inextensibility [Figs. S1D and S2B]. However, we find that an inextensible Rouse polymer still bends in response to activity modulations [Figs. S1D,H and S2B,C], as indicated by the second term of Eq. (S44).

***A.5. Correlated excitations induce folding.*** Now that we have seen how activity modulations lead to the bending of a Rouse polymer, we study a generalized scenario where the athermal excitations are correlated along the polymer backbone. Thus, we consider a non-diagonal correlation function $\mathcal{C}(s,s') = \bar{\xi}^{-1}\,A_0\,[\delta(s-s') + \widehat{\mathcal{C}}(s,s')]$. The spectrum of these correlated excitations is determined by the Fourier transform $\widehat{\mathcal{C}}(s,s') \to \widehat{C}_{qk}$ as defined by Eq. (S4), so that one has $C_{qk} = \bar{\xi}^{-1}\,A_0\,[L\delta_{qk} + \widehat{C}_{qk}]$. As before in section 2 A.3 "Activity modulations lead to bending", we keep the friction coefficient of each material point fixed, $\xi(s) = \bar{\xi}$, and consider homogeneous line tension, $\kappa(s) = \bar{\kappa}$. The response matrix of the polymer is therefore diagonal, $J_{qk} = \bar{\xi}^{-1}\,\bar{\kappa}\,q^2\,\delta_{qk}$, so that Eq. (S28) and Eq. (S20) yield the same result. Taken together, we finally evaluate Eq. (S20) to determine the steady-state polymer conformation:

$$X_{qk} = b^2 \left[ \frac{L\delta_{qk}}{q^2} + \frac{2\,\widehat{C}_{qk}}{q^2 + k^2} \right] , \qquad \text{[S46]}$$

where the characteristic length is given by $b \coloneqq \sqrt{A_0/(2\bar{\kappa})}$, as before.

Taking the limit of an infinitely long polymer, $L \to \infty$ so that $L\delta_{qk} \to 2\pi\,\delta(q-k)$, and transforming Eq. (S46) back into real space quantifies the effect of correlated excitations (through active processes) on the conformation of a Rouse polymer:

$$\langle \boldsymbol{r}(s,t) \cdot \boldsymbol{r}(s',t) \rangle = -\frac{1}{2}\, b^2 \left[ \|s - s'\| + \iint dx\, dx'\, \widehat{\mathcal{C}}(x,x')\, G^A(s-x, s'-x') \right] , \qquad \text{[S47]}$$

where the Green's function kernel is given by Eq. (S42):

$$G^A(s_1, s_2) = \frac{1}{\pi} \left[ 2\gamma_e + \log\left( s_1^2 + s_2^2 \right) \right] . \qquad \text{[S42]}$$

As we have discussed in the main text and will further explain in section 2 B "Passive Rouse polymer with weak long-ranged harmonic interactions", such correlated active processes can fold the Rouse polymer into any desired conformation.

***A.6. Characteristic features of polymer folding driven by correlated excitations.*** In this section, we use our analytical theory to further investigate the shape changes introduced by correlated activity. For a scenario that is analogous to the main text, we consider a block-shaped correlation function. Such a block-shaped correlation function can be decomposed into individual square boxes[5] with widths $\lambda_i$ and center coordinates $(s_i, s_i')$. The vertical edges of each box correspond to the boundary coordinates $s_{i\pm} \coloneqq s_i \pm \lambda_i/2$ of a polymer segment $[s_{i-}, s_{i+}]$ whereas the horizontal edges indicate a (possibly different) polymer segment $[s_{i-}', s_{i+}']$. The athermal excitations in both polymer segments have pairwise correlation coefficient $\rho_i$, which either indicates average motion in the same direction ($\rho_i > 0$) or in opposite directions ($\rho_i < 0$). To ensure that the correlation function is positive definite, boxes on the diagonal ($s_i = s_i'$) have a positive correlation coefficient, whereas each off-diagonal box ($s_i \neq s_i'$) has a symmetric counterpart and introduces two additional boxes on the diagonal.

We measure the shape that the polymer adopts in response to these excitations by the mean scalar product between tangent vectors $\boldsymbol{\tau}(s,t) \coloneqq b^{-1}\,\partial_s \boldsymbol{r}(s,t)$ at different material points[6] [Fig. S3A]. When comparing opposing polymer segment borders, $(s_{i\pm}, s_{i\mp}')$, our theory predicts that correlated excitations ($\rho_i > 0$) will lead to anti-alignment of the respective tangent vectors. In contrast, these tangent vectors will align for anticorrelated excitations ($\rho_i < 0$), which is only possible in off-diagonal boxes that relate different polymer segments ($s_i \neq s_i'$). When comparing matching polymer segment borders, $(s_{i\pm}, s_{i\pm}')$, our theory predicts that correlated excitations ($\rho_i > 0$) will lead to alignment (and stretching when we consider a box on the diagonal) of the respective tangent vectors. In contrast, these tangent vectors will anti-align for anticorrelated excitations ($\rho_i < 0$), which is only possible in off-diagonal boxes that relate different polymer segments ($s_i \neq s_i'$). To cast these results into a geometric picture, in the following we propose polymer shapes that satisfy above criteria. To that end, we use a two-dimensional projection that falls within the scope of our theory.

In an individual polymer segment, correlated excitations drive all material points in the same direction [Fig. S3B, wide arrows and boxes on the diagonal]. This effect will yank the neighboring polymer segments closer together (and stretch their polymer backbone), analogous to an increase in statistically independent activity, and thereby fold the active polymer segment [Fig. S3B]. Meanwhile, the shape of the active segment itself remains largely unaffected [tangent correlation function in Fig. S3A]. Pairs of active polymer segments that typically move in the same direction ($\rho_i > 0$) will align their folds, while polymer segments moving in opposite directions ($\rho_i < 0$) will exhibit opposing folds [Fig. S3B]. One can create more folds and control their relative alignment by having additional active polymer segments, thereby increasing the complexity of the folded polymer. Note that these alignment effects are independent of the separation between the active polymer segments, so that correlated active processes can induce longer-ranged structure in the polymer shape than activity modulations alone. This increase in complexity can be further rationalized by counting the number of independent entries in the correlation matrix of the athermal excitations, for an active polymer made of $N$ monomers. While statistically independent excitations have a diagonal correlation matrix with only $N$ independent entries, correlated excitations are characterized by a non-diagonal correlation matrix with $N(N+1)/2$ independent entries. These additional parameters can then address all of the $N(N-1)/2$ independent entries of the mean squared separation map (whose diagonal entries vanish) and even the $N$ entries of the average squared monomer distance to the origin.

### B. Passive Rouse polymer with weak long-ranged harmonic interactions.

[5]Rectangles can be treated analogously and do not qualitatively change the results.

[6]This tangent autocorrelation function can be determined from the pairwise position correlation function, Eq. (S47).

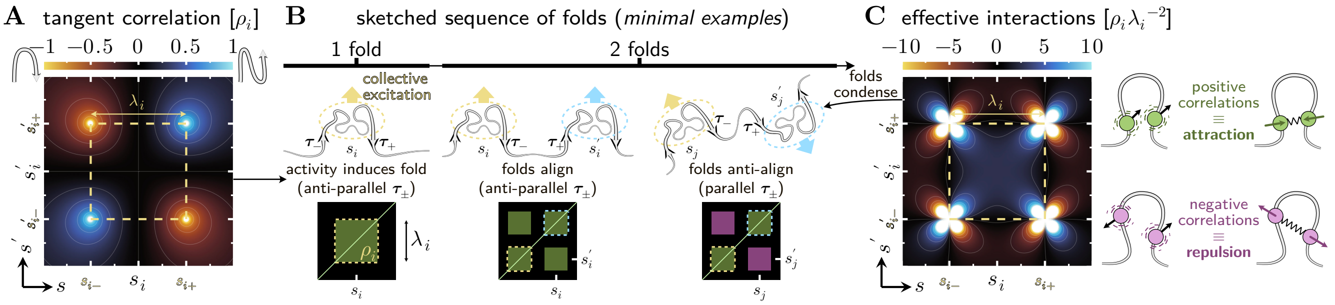

Fig. S3. **Correlated activity can guide polymer folding into specific shapes.** We use our analytical theory to study the effect of correlated athermal excitations on polymer shape, by decomposing any block-structured correlation function into its constituent blocks of width $\lambda_i$. Each box corresponds to pairs of polymer segments that experience correlated excitations, and whose boundaries in sequence space are indicated by orange dashed lines. **A)** Material points at opposing boundaries have anti-aligned tangent vectors (red colors in the colormap) while matching boundaries have aligned tangent vectors (blue colors in the colormap); this trend is inverted for anti-corrrelations ($\rho_i < 0$, thus reversing the colormap). **B)** By using our results from panel A), we propose polymer conformations that have the correct (anti-)alignment of tangent vectors at the boundaries of the correlated regions (dashed ellipse). Below each sketched example, we depict the corresponding correlation function of the excitations $\mathcal{C}(s, s')$. Black shaded arrows indicate the typical direction of the tangent vectors. Correlated excitations of individual polymer segments (box on the diagonal) induce folding, as all material points within the respective segment typically move in the same direction (green arrow) and yank the neighboring segments closer together. Pairs of polymer segments that experiences (anti-)correlated excitations (boxes that are located off-diagonal) will not only fold locally, but also (anti-)align these folds relative to each other. **C)** Identical to Fig. 3D in the main text. Correlated athermal excitations of polymer segments centered around $s_i$ and $s_i'$ lead to an effective attraction through weak, long-ranged harmonic interactions. Conversely, anti-correlated excitations of these segments lead to an effective repulsion. Thus, we expect that individual segments experiencing correlated excitations should adopt a compact shape.

***B.1. Setup and analytical solution of passive polymer model.*** In this section, we construct an effective model where the polymer folds not due to correlated excitations generated by active processes, but because of weak long-ranged harmonic interactions between different material points. In this linearized model, the dynamics of the polymer are determined by

$$\partial_t \boldsymbol{r}(s,t) = \frac{\bar{\kappa}}{\bar{\xi}} \left\{ \partial_s^2 \boldsymbol{r}(s,t) + \int ds' \, \widehat{\mathcal{K}}(s,s') \left[ \boldsymbol{r}(s',t) - \boldsymbol{r}(s,t) \right] \right\} + \boldsymbol{\eta}(s,t) \,, \tag{S48}$$

where $\widehat{\mathcal{K}}(s,s')$ is a dimensionless field that encodes harmonic interactions between different material points, and $\boldsymbol{\eta}(s,t)$ is a Gaussian random displacement velocity field with zero mean. We consider statistically independent excitations with homogeneous magnitude,

$$\langle \boldsymbol{\eta}(s,t) \cdot \boldsymbol{\eta}(s',t') \rangle = \mathcal{C}(s,s') \, \delta(t-t') \,, \quad \text{with} \quad \mathcal{C}(s,s') = \bar{\xi}^{-1} A_0 \, \delta(s-s') \,, \tag{S49}$$

which corresponds to $C_{qk} = \bar{\xi}^{-1} A_0 \, L \delta_{qk}$. Therefore, the polymer will not fold in the absence of harmonic interactions, that is when $\widehat{\mathcal{K}}(s,s') = 0 \, \forall \, s, s'$. Within the formalism introduced in section 1 A "Rouse mode decomposition leads to a non-diagonal system of equations", the response matrix of the polymer is given by a non-diagonal matrix

$$J_{qk} = \bar{\xi}^{-1} \, \bar{\kappa} \, \left[ q^2 \, \delta_{qk} - L^{-1} \left( \widehat{K}_{qk} - \widehat{K}_{q-k,0} \right) \right] \,, \tag{S50}$$

where we have used the following convention for Fourier transforms [cf. Eq. (S4)]:

$$\widehat{K}_{qk} := \iint ds \, ds' \, e^{-iqs} \widehat{\mathcal{K}}(s,s') e^{iks'} \,. \tag{S51}$$

In the response matrix, Eq. (S50), we identify the diagonal entries $\bar{J}_{qk} = \bar{\xi}^{-1}\,\bar{\kappa}\,q^2\,\delta_{qk}$ and the off-diagonal entries $\delta J_{qk} = -\bar{\xi}^{-1}\,\bar{\kappa}\,L^{-1}\,[\widehat{K}_{qk} - \widehat{K}_{q-k,0}]$. Next, we investigate how these harmonic interactions affect the polymer conformation.

In the following, we consider reciprocal harmonic interactions, $\widehat{\mathcal{K}}(s,s') = \widehat{\mathcal{K}}(s',s)$, which means that the response matrix is Hermitian. In this scenario, one can evaluate the formal solution for the conformation of the polymer in Fourier space, Eq. (S16), without relying on a perturbation approach:

$$\boldsymbol{X} = \frac{1}{2}\,\bar{\xi}^{-1}\,A_0\,L\,\boldsymbol{J}^{-1} = \frac{1}{2}\,\bar{\xi}^{-1}\,A_0\,L\,\left(\bar{\boldsymbol{J}} + \delta\boldsymbol{J}\right)^{-1} \approx \frac{1}{2}\,\bar{\xi}^{-1}\,A_0\,L\,\left(\bar{\boldsymbol{J}}^{-1} - \bar{\boldsymbol{J}}^{-1}\cdot\delta\boldsymbol{J}\cdot\bar{\boldsymbol{J}}^{-1}\right)\,, \qquad \text{[S52]}$$

where in the last step we have assumed that the long-ranged harmonic coupling is weak. In other words, we assume that the mechanical properties of the polymer are dominated by line tension, which is the same assumption as the one underlying our perturbation approach in section 1 E "Perturbation approximation for non-diagonal response matrices". Finally, we evaluate Eq. (S52):

$$X_{qk} = b^2\left[\frac{L\delta_{qk}}{q^2} + \frac{\widehat{K}_{qk} - \widehat{K}_{q-k,0}}{q^2\,k^2}\right]\,, \qquad \text{[S53]}$$

where the characteristic length is given by $b \coloneqq \sqrt{A_0/(2\bar{\kappa})}$, as before. Note that the homogeneous Fourier modes of the harmonic interaction map, $q=0$ and $k=0$, do not contribute to the polymer conformation. On a more fundamental level, the homogeneous Fourier modes of the harmonic interaction map cancel out in the polymer's equation of motion, Eq. (S48). Therefore, in the following we assume $\widehat{K}_{q0} = 0\,\forall\,q$, which means that there is no homogeneous global coupling through harmonic interactions. As expected, Eq. (S53) shows that harmonic interactions can fold the polymer into complex shapes. In the following, we refer to this as the "passive model".

***B.2. Comparison to active polymer with correlated excitations.*** We now compare the steady-state conformation of the Rouse polymer in our passive model, Eq. (S53), to the steady-state conformation of a Rouse polymer without long-ranged harmonic interactions, but instead subject to correlated athermal excitations ("active model"), cf. Eq. (S46). In section 2 A.5 "Correlated excitations induce folding", we discussed that the active model has a diagonal response matrix given by $\bar{J}_{qk}$, which is identical to the diagonal entries of the response matrix in the passive model, Eq. (S50). In contrast to the passive model, however, the active model is subject to athermal excitations characterized by a non-diagonal correlation function $\mathcal{C}(s,s') = \bar{\xi}^{-1}\,A_0\,[\delta(s-s') + \widehat{\mathcal{C}}(s,s')]$. The spectrum of these correlated excitations is determined by the Fourier transform $\widehat{\mathcal{C}}(s,s') \to \widehat{C}_{qk}$ as defined by Eq. (S4), so that one has $C_{qk} = \bar{\xi}^{-1}\,A_0\,[L\delta_{qk} + \widehat{C}_{qk}]$. These two fundamentally different models will result in the same steady-state polymer conformation, if the following criterion is satisfied:

$$\widehat{K}_{qk} = \widehat{C}_{qk}\,\frac{2\,q^2\,k^2}{q^2 + k^2}\,. \qquad \text{[S54]}$$

Taking the limit of an infinitely long polymer, $L \to \infty$ so that $L\delta_{qk} \to 2\pi\,\delta(q-k)$, and transforming Eq. (S54) back into real space shows that correlated athermal excitations induce effective long-ranged harmonic interactions between different material points,

$$\widehat{\mathcal{K}}(s,s') = -\left[\partial_s^2 + \partial_{s'}^2\right]\,\widehat{\mathcal{C}}(s,s') + \iint dx\,dx'\,\widehat{\mathcal{C}}(x,x')\,G^K(s-x,s'-x')\,, \qquad \text{[S55]}$$

with the following Green's function kernel:

$$G^K(s_1, s_2) = -\frac{6}{\pi}\,\frac{s_1^4 + s_2^4 - 6s_1^2 s_2^2}{(s_1^2+s_2^2)^4}\,. \qquad \text{[S56]}$$

In Eq. (S55), the first term indicates that pairs of material points whose athermal excitations are maximally correlated will attract each other, whereas maximally anticorrelated excitations lead to repulsion. The second term in Eq. (S55) indicates that pairs of material points whose athermal excitations are correlated (anticorrelated) in general will attract (repel) each other [Fig. S3C], as discussed in the main text. Note that the Green's kernel has the most convenient representation in polar coordinates $G^K(\alpha\cos\phi, \alpha\sin\phi) = -6\cos(4\phi)/(\pi\alpha^4)$. This shows that the convolution term in Eq. (S55) will vanish if the correlation function of the athermal excitations, $\widehat{\mathcal{C}}(s,s')$, is rotationally symmetric around some point $(s,s')$ in the sequence correlation space. In that case, only the first term in Eq. (S55) will remain.

***B.3. Comparison to active polymer with activity modulations.*** We now study the effective interactions that arise in an active Rouse polymer that is subject to statistically independent athermal excitations with modulations in magnitude, $\mathcal{C}(s,s') = \bar{\xi}^{-1} A_0\,[1+\epsilon(s)]\,\delta(s-s')$. The spectrum of these activity modulations is determined by the Fourier transform $\epsilon(s) \to \epsilon_q$ as defined by Eq. (S3), so that one has $C_{qk} = \bar{\xi}^{-1} A_0\,[L\delta_{qk} + \epsilon_{q-k}]$ where $\epsilon_0 = 0$. Comparing with the previous section, one has $\widehat{C}_{qk} \equiv \epsilon_{q-k}$ in the equivalence criterion, Eq. (S54). Taking the limit of an infinitely long polymer, $L \to \infty$ so that $L\delta_{qk} \to 2\pi\,\delta(q-k)$, and transforming Eq. (S54) back into real space yields:

$$\widehat{\mathcal{K}}(s,s') = \frac{3}{4}\,\epsilon''\!\left(\frac{s+s'}{2}\right)\delta(s-s') - \epsilon\!\left(\frac{s+s'}{2}\right)\delta''(s-s') + \int\frac{dq}{2\pi}\left[\epsilon_q\,\frac{\|q\|^3}{4}\,e^{-\frac{\|q\|}{2}\,\|s-s'\|}\right]e^{iq\frac{s+s'}{2}}\,, \qquad \text{[S57]}$$

where $\epsilon''(s) \coloneqq \partial_s^2\epsilon(s)$ and $\delta''(s) \coloneqq \partial_s^2\delta(s)$ are second derivatives. To understand the effect of the harmonic interaction map, Eq. (S57), we substitute it into the corresponding term of the equation of motion, Eq. (S48):

$$\int ds'\,\widehat{\mathcal{K}}(s,s')\,[\boldsymbol{r}(s') - \boldsymbol{r}(s)] = -\partial_s\Big[\epsilon(s)\,\partial_s\boldsymbol{r}(s)\Big] + \int ds'\,\widehat{\mathcal{K}}_{\mathrm{NL}}(s,s')\,[\boldsymbol{r}(s') - \boldsymbol{r}(s)]\,, \qquad \text{[S58]}$$

where we have defined the map of *non-local* interactions

$$\widehat{\mathcal{K}}_{\mathrm{NL}}(s,s') = \int\frac{dq}{2\pi}\left[\epsilon_q\,\frac{\|q\|^3}{4}\,e^{-\frac{\|q\|}{2}\,\|s-s'\|}\right]e^{iq\frac{s+s'}{2}}\,. \qquad \text{[S59]}$$

In addition to these non-local interactions, the first term on the right-hand side of Eq. (S58) shows that modulations in activity also induce an effective decrease (increase) of line tension in more (less) active regions [Fig. S4B, central panel]. These changes in effective line tension are responsible for the expansion or contraction of the polymer backbone, respectively, which we observed. Consistent with these findings, making the polymer inextensible as outlined in section 2 A.4 "Enforcing inextensibility in active polymers" will cancel out the modulations in effective line tension, and only leave non-local interactions.

In contrast, the non-local interactions, Eq. (S59), explain activity-induced polymer bending and straightening. Non-neighboring material points in regions with increased activity show an effective pairwise attraction [Fig. S4B], which leads to bending [Fig. S4A]. Conversely, polymer straightening [Fig. S4A] is associated with an effective pairwise repulsion between non-neighboring material points

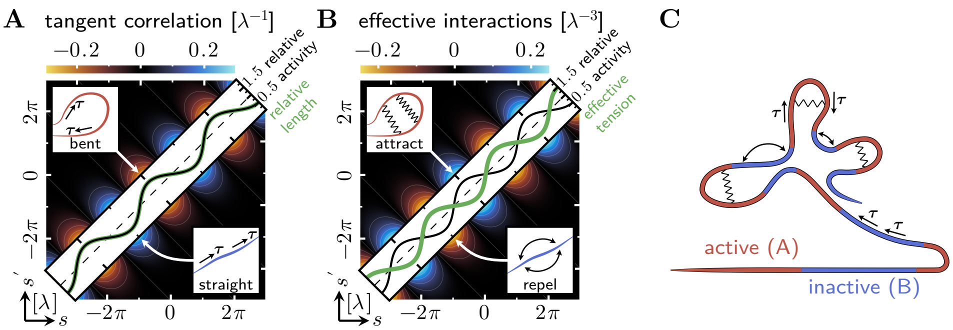

Fig. S4. **Activity modulations induce polymer folding. A)** Correlation between tangent vectors at different material points. A local increase in activity with a characteristic length scale $\lambda$ leads to polymer bending (anti-correlations), while a local decrease in activity leads to polymer straightening (positive correlations). **B)** An active polymer driven by activity modulations folds into the same steady-state conformation as a passive polymer with homogeneous activity $A_0$ and long-ranged pairwise harmonic interactions, cf. Eq. (S55). This mapping explains that a local increase in activity leads to expansion of the polymer backbone through an effective decrease in line tension (diagonal panel). Conversely, a local decrease in activity leads to condensation of the polymer backbone through an effective increase in line tension (diagonal panel). Polymer bending is induced by an effective attraction between material points that are not nearest neighbors but lie within the same polymer segment (annotation in top left panel), while polymer straightening is mediated by an effective repulsion (annotation in bottom right panel). **C)** Activity modulations fold the polymer into a "flower" shape, where active regions loop around straighter inactive segments.

in regions with decreased activity [Fig. S4B]. Thus, local activity modulations can be mapped to harmonic interactions between distal material points. To conclude, we have here demonstrated that the steady state shape of an active polymer can be directly mapped to a passive surrogate model [Fig. S4C].

### 3. Subdiffusive dynamics of active polymers

**A. Mean squared displacement of a locus in an active polymer.** To extract characteristic dynamic features of active polymers, we will now determine the mean squared distance $\langle[\boldsymbol{r}(s,t+\Delta t)-\boldsymbol{r}(s,t)]^2\rangle$ that a specific locus travels within the time window $\Delta t$. To that end, we specialize the derivation outlined in section 1 F "Mean squared distance traveled by a specific polymer locus". As before in section 1 D "Conformation of a polymer with a diagonal response matrix", we consider a polymer that has a diagonal response matrix, so that Eq. (S31) evaluates to

$$\Big\langle[\boldsymbol{r}(s,t+\Delta t)-\boldsymbol{r}(s,t)]^2\Big\rangle = \sum_{qk} e^{i(q-k)s}\Bigg\{\Big(1-e^{-J_{qq}\Delta t}\Big)\frac{C_0\,\delta_{qk}}{J_{qq}} + \Big(1-e^{-J_{qq}\Delta t}\Big)\frac{2C_0\,\delta\widehat{C}_{qk}}{J_{qq}+J_{kk}}\Bigg\}\,. \tag{S60}$$

More specifically, the response matrix of the Rouse polymer is given by $J_{qk}=\bar{\xi}^{-1}\,\bar{\kappa}\,q^2\,\delta_{qk}$, with homogeneous friction coefficient $\bar{\xi}$ and line tension $\bar{\kappa}$. For the athermal excitations that drive the polymer's motion, we again consider the correlation function $\mathcal{C}(s,s')=\bar{\xi}^{-1}\,A_0\,[\delta(s-s')+\widehat{\mathcal{C}}(s,s')]$, so that one has $C_{qk}=\bar{\xi}^{-1}\,A_0\,[L\delta_{qk}+\widehat{C}_{qk}]$. A comparison with Eq. (S26) suggests $C_0=\bar{\xi}^{-1}\,A_0\,L$ and $\delta\widehat{C}_{qk}=L^{-1}\widehat{C}_{qk}$, which we substitute into Eq. (S60). Taking the limit of an infinitely long polymer, $L\to\infty$ so that $L\delta_{qk}\to 2\pi\,\delta(q-k)$, and representing the elapsed time $\Delta t=\bar{\xi}\bar{\kappa}^{-1}\,\tau$ by a dimensionless variable $\tau$, we arrive at

$$\Big\langle[\boldsymbol{r}(s,t+\bar{\xi}\bar{\kappa}^{-1}\,\tau)-\boldsymbol{r}(s,t)]^2\Big\rangle = 2b^2\left[\sqrt{\frac{\tau}{\pi}}+2\int\frac{dq}{2\pi}\int\frac{dk}{2\pi}\,e^{i(q-k)s}\,\widehat{C}_{qk}\,\frac{1-e^{-q^2\tau}}{q^2+k^2}\right]\,, \tag{S61}$$

where the characteristic length is given by $b:=\sqrt{A_0/(2\bar{\kappa})}$, as before. Using this result, we can predict the transport properties of individual loci on active polymers that are driven by activity modulations, or correlated excitations.

Equation (S61) can be simplified if the active polymer is driven only by statistically independent excitations, characterized by $\widehat{\mathcal{C}}(s,s')\equiv\epsilon(s)\,\delta(s-s')$, with inhomogeneous activity $A(s)=A_0\,[1+\epsilon(s)]$, so that $\widehat{C}_{qk}\equiv\epsilon_{q-k}$. Then, one has

$$\Big\langle[\boldsymbol{r}(s,t+\bar{\xi}\bar{\kappa}^{-1}\,\tau)-\boldsymbol{r}(s,t)]^2\Big\rangle = 2b^2\sqrt{\frac{\tau}{\pi}}\left[1+2\sqrt{\pi}\int\frac{dk}{2\pi}\,\epsilon_k\,e^{iks}\int\frac{dq}{2\pi}\,\frac{1-e^{-q^2}}{q^2+(q-\sqrt{\tau}\,k)^2}\right]\,. \tag{S62}$$

This representation reveals that for $\tau\ll 1$ the mean squared traveled distance grows as

$$\Big\langle[\boldsymbol{r}(s,t+\bar{\xi}\bar{\kappa}^{-1}\,\tau)-\boldsymbol{r}(s,t)]^2\Big\rangle = 2b^2\sqrt{\frac{\tau}{\pi}}\,[1+\epsilon(s)]\,. \tag{S63}$$

Thus, a local increase in activity enhances the subdiffusive motion of the affected loci, whereas a local decrease in activity reduces this subdiffusive motion [Fig. S5A]. On sufficiently long timescales, $\tau\to\infty$, the subdiffusive motion of every locus is determined by the mean level of activity [Fig. S5A]. For sinusoidal activity modulations, i.e. $\widehat{\mathcal{C}}(s,s')\equiv\epsilon\,\cos(s/\lambda)\,\delta(s-s')$ so that $\widehat{C}_{qk}\equiv(\epsilon/2)\,[2\pi\delta(q-k-\lambda^{-1})+2\pi\delta(q-k+\lambda^{-1})]$, Eq. (S61) is given by

$$\Big\langle[\boldsymbol{r}(s,t+\bar{\xi}\bar{\kappa}^{-1}\,\tau)-\boldsymbol{r}(s,t)]^2\Big\rangle = 2\lambda\,b^2\left[\sqrt{\frac{\tau/\lambda^2}{\pi}}+\epsilon\,\cos(s/\lambda)\,g_{\mathrm{a}}^{\epsilon}(\tau/\lambda^2)\right]\,, \tag{S64a}$$

with

$$g_{\mathrm{a}}^{\epsilon}(\tau)=\int\frac{dq}{\pi}\,\frac{1-e^{-q^2\tau}}{q^2+(q+1)^2}\,, \tag{S64b}$$

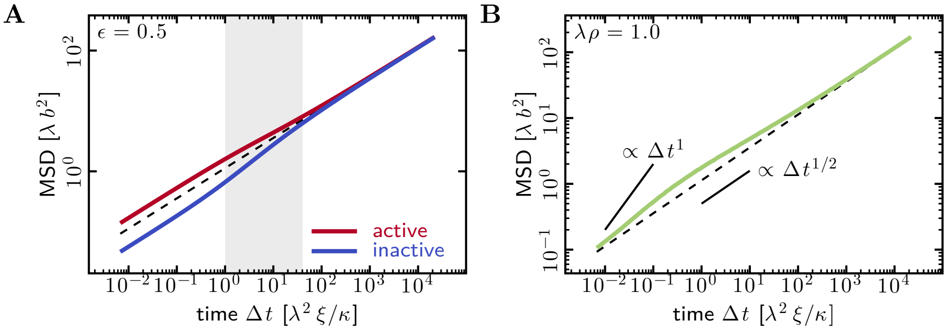

Fig. S5. **Mean squared displacement of a specific locus in an active polymer in a given time $\Delta t$. Dashed lines show dynamics for a homogeneous reference polymer. A)** The active polymer is driven by statistically independent excitations with correlation function $\mathcal{C}(s,s') \equiv \bar{\xi}^{-1}\, A(s)\, \delta(s-s')$ and sinusoidal activity modulations $A(s) = A_0\, [1 + \epsilon\, \cos(s/\lambda)]$. On short time scales, the local level of activity determines the subdiffusive motion of every locus, with a characteristic $\Delta t^{1/2}$-scaling. On long time scales, the subdiffusive motion of every locus is determined by the average level of activity of the polymer. Note that, if one were to only measure the mean squared displacement in the transition window (gray), then one would observe different subdiffusion exponents for the active and the inactive regions. This effect depends on the amplitude of the activity modulations, $\epsilon$, and vanishes for $\epsilon \to 0$. **B)** The active polymer is driven by coherent excitations. Specifically, every material point within a contiguous segment of length $\lambda$ experiences excitations with correlation $\mathcal{C}(s,s') = \bar{\xi}^{-1}\, A_0\, [\delta(s-s') + \widehat{\mathcal{C}}(s,s')]$ and $\widehat{\mathcal{C}}(s,s') = \rho\ \forall\ s,s' \in [s_0 - \lambda/2, s_0 + \lambda/2]$ else zero. The motion of the material point at the center of this segment, $s = s_0$, shows a characteristic $\Delta t^{1/2}$-scaling both on short and on long time scales. At intermediate times, there is an inhomogeneous contribution with a characteristic $\Delta t^1$-scaling.

which we solve by numerical integration.

Next, we consider an active polymer that is driven by correlated excitations. Specifically, we consider a scenario where every material point within a contiguous segment of length $\lambda$ experiences excitations with correlation $\mathcal{C}(s,s') = \bar{\xi}^{-1}\, A_0\, [\delta(s-s') + \widehat{\mathcal{C}}(s,s')]$ and $\widehat{\mathcal{C}}(s,s') = \rho\ \forall\ s,s' \in [s_0 - \lambda/2, s_0 + \lambda/2]$ else zero. Then, Eq. (S61) is given by

$$\left\langle [\boldsymbol{r}(s_0, t + \bar{\xi}\bar{\kappa}^{-1}\,\tau) - \boldsymbol{r}(s_0,t)]^2 \right\rangle = 2\lambda\, b^2 \left[ \sqrt{\frac{\tau/\lambda^2}{\pi}} + \rho\,\lambda\, g_{\mathrm{a}}^{\rho}(\tau/\lambda^2) \right] , \qquad \text{[S65a]}$$

with

$$g_{\mathrm{a}}^{\rho}(\tau) = 2 \int \frac{dq}{\pi} \int \frac{dk}{\pi}\, \frac{\sin(q/2)}{q}\, \frac{\sin(k/2)}{k}\, \frac{1 - e^{-q^2\tau}}{q^2 + k^2} , \qquad \text{[S65b]}$$

which we solve by numerical integration. In contrast to activity modulations where the mean squared traveled distance scales characteristically $\propto \tau^{1/2}$, correlated excitations induce an inhomogeneous contribution with a different scaling $\propto \tau^1$ [Fig. S5B]. This can be intuitively understood by considering the limiting case where every material point of a contiguous polymer segment experiences the same random force (i.e., the excitations are perfectly correlated). Then, the entire polymer segment will show coherent diffusion equivalent to a single large particle, which leads to the $\tau^1$-scaling.

**B. Mean squared displacement of a locus in a passive polymer.** To extract characteristic dynamic features of passive polymers, we will now determine the mean squared distance $\langle [\boldsymbol{r}(s, t{+}\Delta t) - \boldsymbol{r}(s,t)]^2 \rangle$ that a specific locus travels within the time window $\Delta t$. To that end, we specialize the derivation outlined in section 1 F "Mean squared distance traveled by a specific polymer locus". We consider a Rouse polymer that is driven by homogeneous excitations, see Eq. (S49), so that $C_{qk} = \bar{\xi}^{-1}\, A_0\, L\delta_{qk}$

and Eq. (S31) evaluates to

$$\left\langle [\boldsymbol{r}(s,t+\Delta t)-\boldsymbol{r}(s,t)]^2 \right\rangle = \sum_{qk} e^{i(q-k)s} \left\{ \left(1-e^{-J_{qq}\Delta t}\right) \frac{C_0\,\delta_{qk}}{J_{qq}} - \frac{C_0\,\delta J_{qk}}{J_{qq}J_{kk}} \left[1-\frac{J_{qq}e^{-J_{kk}\Delta t}-J_{kk}e^{-J_{qq}\Delta t}}{J_{qq}-J_{kk}}\right] \right\}. \quad \text{[S66]}$$

As discussed in section 2 B.2 "Comparison to active polymer with correlated excitations", we compare the dynamics of this passive polymer to an active polymer whose athermal excitations are characterized by the correlation function $\mathcal{C}(s,s') = \bar{\xi}^{-1} A_0 \,[\delta(s-s') + \widehat{\mathcal{C}}(s,s')]$, so that $C_{qk} = \bar{\xi}^{-1} A_0\,[L\delta_{qk} + \widehat{C}_{qk}]$. A comparison with Eq. (S26) suggests $C_0 = \bar{\xi}^{-1} A_0\, L$ and $\delta\widehat{C}_{qk} = L^{-1}\widehat{C}_{qk}$, which we will later substitute. To show the same distribution of conformations in steady state, the passive polymer requires off-diagonal entries in its response matrix, which we assume to be Hermitian and which we derive from Eq. (S28):

$$\frac{\delta J_{qk}}{J_{qq}J_{kk}} = -2\frac{\delta\widehat{C}_{qk}}{J_{qq}+J_{kk}}\,. \quad \text{[S67]}$$

For a Rouse polymer, the diagonal elements of the response matrix are given by $J_{qq} = \bar{\xi}^{-1}\,\bar{\kappa}\, q^2$, with homogeneous friction coefficient $\bar{\xi}$ and line tension $\bar{\kappa}$. We substitute the diagonal elements of the response matrix, as well as its off-diagonal elements Eq. (S67), into Eq. (S66). Taking the limit of an infinitely long polymer, $L\to\infty$ so that $L\delta_{qk} \to 2\pi\,\delta(q-k)$, and representing the elapsed time $\Delta t = \bar{\xi}\bar{\kappa}^{-1}\,\tau$ by a dimensionless parameter $\tau$, we arrive at

$$\left\langle [\boldsymbol{r}(s,t+\bar{\xi}\bar{\kappa}^{-1}\,\tau)-\boldsymbol{r}(s,t)]^2 \right\rangle = 2b^2 \left[ \sqrt{\frac{\tau}{\pi}} + 2\int\frac{dq}{2\pi}\int\frac{dk}{2\pi}\, e^{i(q-k)s}\,\widehat{C}_{qk}\, \frac{q^2(1-e^{-k^2\tau})-k^2(1-e^{-q^2\tau})}{q^4-k^4} \right], \quad \text{[S68]}$$

where the characteristic length is given by $b := \sqrt{A_0/(2\bar{\kappa})}$, as before. Using this result, we can predict the transport properties of individual loci on passive polymers.

To compare with the results presented in section 3 A "Mean squared displacement of a locus in an active polymer", we again consider two scenarios. For sinusoidal activity modulations, i.e. $\widehat{\mathcal{C}}(s,s') \equiv \epsilon\,\cos(s/\lambda)\,\delta(s-s')$ so that $\widehat{C}_{qk} \equiv (\epsilon/2)\,[2\pi\delta(q-k-\lambda^{-1}) + 2\pi\delta(q-k+\lambda^{-1})]$, Eq. (S68) is given by

$$\left\langle [\boldsymbol{r}(s,t+\bar{\xi}\bar{\kappa}^{-1}\,\tau)-\boldsymbol{r}(s,t)]^2 \right\rangle = 2\lambda\, b^2 \left[ \sqrt{\frac{\tau/\lambda^2}{\pi}} + \epsilon\,\cos(s/\lambda)\, g_{\mathrm{p}}^{\epsilon}(\tau/\lambda^2) \right], \quad \text{[S69a]}$$

with

$$g_{\mathrm{p}}^{\epsilon}(\tau) = \int\frac{dq}{\pi}\, \frac{q^2\,(1-e^{-(q+1)^2\tau}) - (q+1)^2\,(1-e^{-q^2\tau})}{q^4-(q+1)^4}\,, \quad \text{[S69b]}$$

which we solve by numerical integration. Note that Eq. (S69) only differs from Eq. (S64) by the inhomogeneous contributions $g_{\mathrm{a/p}}^{\epsilon}(\tau)$. Next, we consider an active polymer that is driven by correlated excitations. Specifically, we consider a scenario where every material point within a contiguous

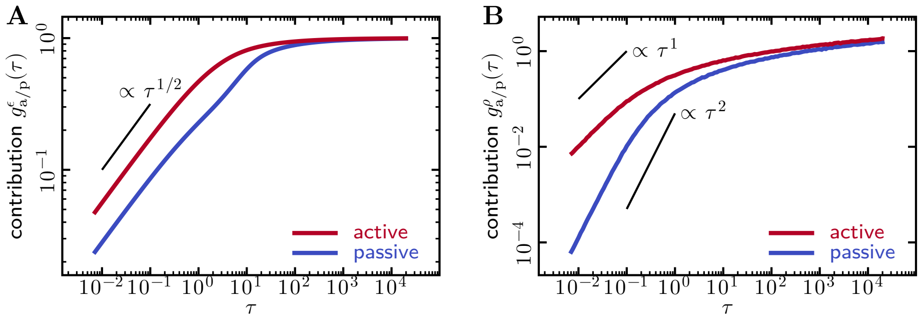

Fig. S6. **Inhomogeneous contributions to mean squared displacement (MSD) that characterize how the dynamics of inhomogeneous active/passive polymers deviate from the dynamics of a homogeneous reference polymer. A)** Inhomogeneous contribution to the MSD of a locus in an active polymer driven by activity modulations [cf. Eq. (S64b)], or a passive polymer [cf. Eq. (S69b)] that reproduces the same distribution of conformations. For the active polymer, we consider statistically independent excitations with correlation function $\mathcal{C}(s,s') \equiv \bar{\xi}^{-1}\,A(s)\,\delta(s-s')$ and sinusoidal activity modulations $A(s) = A_0\,[1+\epsilon\,\cos(s/\lambda)]$. **B)** Inhomogeneous contribution to the MSD for an active polymer driven by correlated excitations [cf. Eq. (S65b)], or a passive polymer [cf. Eq. (S70b)] that reproduces the same distribution of conformations. For the active polymer, every material point within a contiguous segment of length $\lambda$ experiences excitations with correlation $\mathcal{C}(s,s') = \bar{\xi}^{-1}\,A_0\,[\delta(s-s') + \widehat{\mathcal{C}}(s,s')]$ and $\widehat{\mathcal{C}}(s,s') = \rho\ \forall\ s,s' \in [s_0-\lambda/2, s_0+\lambda/2]$ else zero. We track the motion of the material point at the center of this segment, $s = s_0$.

segment of length $\lambda$ experiences excitations with correlation $\mathcal{C}(s,s') = \bar{\xi}^{-1}\,A_0\,[\delta(s-s') + \widehat{\mathcal{C}}(s,s')]$ and $\widehat{\mathcal{C}}(s,s') = \rho\ \forall\ s,s' \in [s_0-\lambda/2, s_0+\lambda/2]$ else zero. Then, Eq. (S68) is given by

$$\left\langle [\boldsymbol{r}(s_0, t+\bar{\xi}\bar{\kappa}^{-1}\,\tau) - \boldsymbol{r}(s_0,t)]^2 \right\rangle = 2\lambda\, b^2 \left[ \sqrt{\frac{\tau/\lambda^2}{\pi}} + \rho\,\lambda\, g^{\rho}_{\mathrm{p}}(\tau/\lambda^2) \right] , \qquad \text{[S70a]}$$

with

$$g^{\rho}_{\mathrm{p}}(\tau) = 2\int \frac{dq}{\pi} \int \frac{dk}{\pi}\, \frac{\sin(q/2)}{q}\, \frac{\sin(k/2)}{k}\, \frac{q^2\,(1-e^{-k^2\tau}) - k^2\,(1-e^{-q^2\tau})}{q^4-k^4} , \qquad \text{[S70b]}$$

which we solve by numerical integration. Note that, again, Eq. (S70) only differs from Eq. (S65) by the inhomogeneous contributions $g^{\rho}_{\mathrm{a/p}}(\tau)$. We find, in general, that active polymers show faster motion than their passive counterparts [Fig. S6].

**C. Monomer mean squared displacement in discrete, active chains.** Having studied the subdiffusive dynamics of active polymers using our continuum theory, we now explore the dynamics of discrete, 1000-mer chains subject to nonlinear constraints. To that end, we use the activity profiles shown in Fig. 2D in the main text, where each monomer is assigned an activity $A_{\mathrm{A}}$ or $A_{\mathrm{B}}$ based on the A/B identities in the Hi-C data from Ref. (8) (Methods). In Fig. S7, we use the activity ratio $A_{\mathrm{A}}/A_{\mathrm{B}} = 5.974$ which produces simulated contact maps that match the compartment score observed in the experimental contact maps of Ref. (8) (see main text). To understand how self avoidance and a spherical confinement affect the dynamics of an active polymer, we first study a discrete Rouse chain using Eq. (S31) with a discrete response matrix; cf. section 1 A.1 "Response matrix of a discrete Rouse polymer depends on chain topology".

In a phantom Rouse chain composed of 1000 monomers (Fig. S7A), the mean squared displacement (MSD) averaged over all Kuhn segments with the same activity, $\mathrm{MSD}(\Delta t) = D^{\mathrm{app}} t^{\alpha}$, shows three regimes. At short time scales, monomers diffuse freely ($\alpha = 1$) with $D_{\mathrm{A}} = A_{\mathrm{A}}/(6\xi)$ and $D_{\mathrm{B}} = A_{\mathrm{B}}/(6\xi)$. At intermediate times, monomers feel the effects of the harmonic springs connecting them to the rest of the chain, producing subdiffusive dynamics where $\alpha = 0.5$. Similar to an infinite, continuous

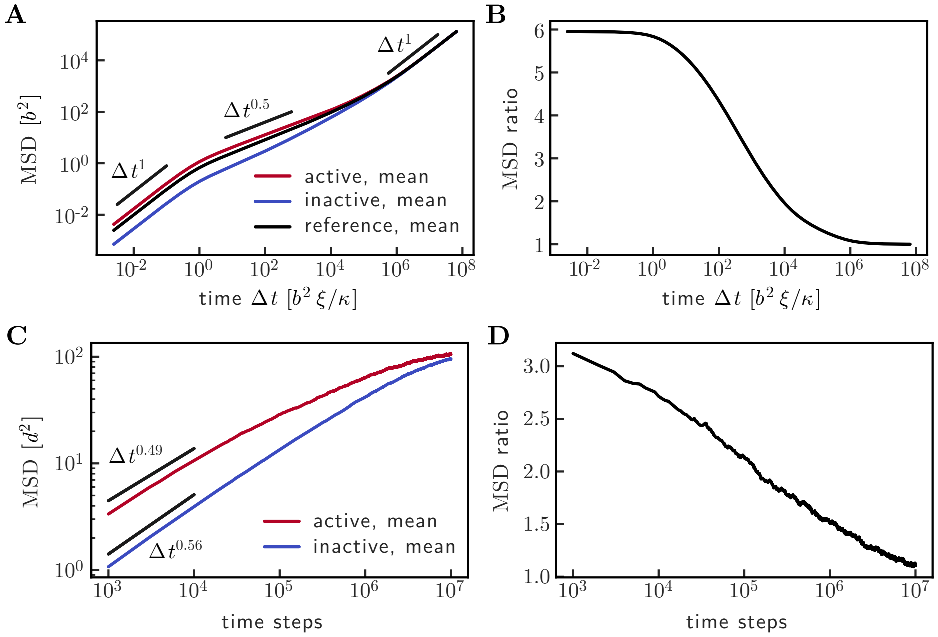

Fig. S7. **Mean squared displacement averaged over active and inactive regions for discrete, 1000-mer chains with activity ratio $A_\mathsf{A}/A_\mathsf{B} = 5.974$.** **A)** For a discrete Rouse chain, the monomer MSD ($\propto \Delta t^\alpha$) of a homogeneous reference polymer (black) displays three regimes: free diffusion of monomers at short times, subdiffusion at intermediate times, and center of mass diffusion at long times. The MSD averaged over active regions is systematically higher and displays a slightly different value of $\alpha$ than that of inactive regions. **B)** During the initial diffusive regime and at the beginning of the subdiffusive regime, the ratio of active and inactive MSDs matches the activity ratio. At long times, the MSD transitions to center-of-mass diffusion, and the MSD ratio approaches $1$. **C)** Ensemble-averaged monomer MSD, in units of monomer diameter squared ($d^2$), computed from $200$ independent simulations of a 1000-mer self avoiding chain in a spherical confinement with volume fraction $0.117$ (same parameters as in Fig. 2D of the main text). The active MSD shows $\alpha < 0.5$, whereas the inactive MSD shows $\alpha > 0.5$. Both curves eventually plateau at the squared radius of confinement. **D)** The ratio of MSDs in active and inactive regions is smaller than the activity ratio during the subdiffusive regime, and eventually approaches $1.0$.

chain, the value of $\alpha$ continuously changes in the subdiffusive regime because both curves eventually coincide. For a discrete chain, this long-time behavior corresponds to the free diffusion of the center of mass of the chain. The MSD of inactive monomers approaches the homogeneous MSD from below, so that $\alpha > 0.5$. Conversely, the MSD of active monomers approaches the homogeneous MSD from above, so that $\alpha < 0.5$. Thus, depending on the time scale of observation and the activity ratio, different values of $\alpha$ can be observed in active and in inactive regions.

The addition of self avoidance, a non-zero spring rest length, and a spherical confinement modifies these dynamics in subtle ways. As seen in Fig. S7C, the active and inactive MSDs are initially subdiffusive and eventually plateau at the squared radius of the confinement. In contrast to the phantom chain, the ratio of MSDs in active and in inactive regions is at least two-fold smaller than the activity ratio during the entirety of the subdiffusive regime [Fig. S7D]. However, the exact value of the MSD ratio and of $\alpha$ in active and inactive regions depends on the volume fraction, the activity ratio, and other microscopic parameters. Thus, even large activity differences could produce small MSD ratios.

### 4. Data-driven inference of the polymer mechanics and the excitations

**A. Inference of the response matrix.** For a polymer with known mechanical properties, which are encoded by the response matrix $\boldsymbol{J}$, our theory establishes a bijective mapping [cf. Eq. (S16)] between athermal excitations with correlation function $\mathcal{C}(s,s')$ and the resulting steady-state conformation. We assume that the polymer has homogeneous mechanical properties and therefore a diagonal response matrix. Therefore, the mean squared separation between different material points is given by [cf. Eq. (S20)]

$$\Delta R^2(s,s') := \left\langle [\boldsymbol{r}(s,t)-\boldsymbol{r}(s',t)]^2 \right\rangle = \frac{1}{L^2}\sum_{qk} \frac{C_{qk}}{J_{qq}+J_{kk}} \left[ e^{i(q-k)s} + e^{i(q-k)s'} - 2e^{iqs-iks'} \right] . \qquad \text{[S71]}$$

If we know the response matrix $\boldsymbol{J}$, then we can now invert the problem and seek the unique correlation function $\mathcal{C}(s,s')$ between the athermal excitations, which fold the polymer into a desired conformation. But what if the mechanical properties of the polymer are not known *a priori*? This lack of information introduces additional modeling degrees of freedom that one needs to constrain based on the provided data and our theory. To that end, we first transform into a coordinate system that is more convenient for the following calculations, $\bar{s} := (s+s')/2$ and $\Delta s := s - s'$. Next, we aim to integrate out inhomogeneities in the mean squared separation map,

$$\overline{\Delta R^2}(\Delta s) := \frac{1}{L-\Delta s}\int_{-\frac{L-\Delta s}{2}}^{\frac{L-\Delta s}{2}} d\bar{s}\, \Delta R^2(\bar{s}+\Delta s/2, \bar{s}-\Delta s/2)\,, \quad \text{with} \quad \Delta s \in [0,L]\,. \qquad \text{[S72]}$$

By making the approximation of a very long polymer ($L \gg \Delta s$), or by considering a polymer whose mean squared separation map is (nearly) translationally invariant, one then has

$$\overline{\Delta R^2}(\Delta s) \approx \frac{1}{L^2}\sum_q \frac{C_{qq}}{J_{qq}} \left[ 1 - e^{iq\Delta s} \right] . \qquad \text{[S73]}$$

To extract the components of the diagonal response matrix, we use a Fourier transform:

$$\int_0^L ds\, e^{-iks}\, \overline{\Delta R^2}(s) = \left[ \frac{1}{L}\sum_q \frac{C_{qq}}{J_{qq}} \right] \delta_{k0} - \frac{1}{L}\frac{C_{kk}}{J_{kk}} . \qquad \text{[S74]}$$

Now, we consider a scenario where translational invariance is broken by athermal excitations with a non-diagonal correlation function $\mathcal{C}(s,s') = \xi^{-1} A_0 [\delta(s-s') + \widehat{\mathcal{C}}(s,s')]$. The spectrum of these correlated excitations is determined by the Fourier transform $\widehat{\mathcal{C}}(s,s') \to \widehat{C}_{qk}$ as defined by Eq. (S4), so that one has $C_{qk} = \xi^{-1} A_0 [L\delta_{qk} + \widehat{C}_{qk}]$. Finally, we make another drastic approximation by assuming that the athermal excitations have no translationally invariant component[7], so that $\widehat{C}_{qq} = 0 \,\forall\, q$. In summary, the response matrix is given by[8]

$$\frac{\xi\, J_{qq}}{A_0} = -\left[ \int_0^L ds\, e^{-iqs}\, \overline{\Delta R^2}(s) \right]^{-1} \quad \forall \quad q \neq 0\,, \qquad \text{[S75]}$$

**B. Heuristic response matrix approximates mechanical properties of simulated nonlinear polymer.** To set up a common model for the chain mechanics, we use a polymer with uniform activity

[7] This assumption excludes all correlation functions that have a contribution $\widehat{\mathcal{C}}(s-s')$.
[8] For discrete polymers, this relation involves discrete spectral transforms.

where Eq. (S75) is exact. We determine the model parameters by fitting the translationally invariant mean squared separation $\Delta R^2(s,s') = \overline{\Delta R^2}(s-s')$ to our artificial data. To that end, we use the following heuristic response matrix[9] [cf. Eq. (S8)]:

$$\frac{\xi\, J_{qq}}{A_0} = \frac{1}{2b^2}\left[\alpha + \left|2\sin\left(\frac{qa}{2}\right)\right|^2 + \kappa_3 \left|2\sin\left(\frac{qa}{2}\right)\right|^3\right], \quad \text{[S76]}$$

with three fit parameters $(b^2, \alpha, \kappa_3)$. The cubic term in Eq. (S76) has only the purpose of improving the fit[10]. We confirm visually that the thus obtained response matrix indeed gives a good approximation of our data [Fig. S8A]. Furthermore, we compare the response matrix obtained through our fitting procedure, Eq. (S76), against the approximated response matrix of simulated polymers that are driven by inhomogeneous activity Eq. (S75). As expected, we find that the inferred mechanical properties of a polymer do not significantly depend on its level of activity [Fig. S8A], which justifies our choice of the homogeneous polymer as reference.

**C. Inference of an activity profile that folds the polymer towards a desired conformation.** Finally, we discuss the details of the optimization procedure used in the main text, which proposes an activity profile that could fold an active polymer as close as possible towards a desired or observed steady-state conformation. We consider a discrete polymer that is driven by statistically independent excitations with covariance $\langle \boldsymbol{\eta}_i(t)\cdot\boldsymbol{\eta}_j(t')\rangle = \xi^{-1} A_i\,\delta_{ij}\,\delta(t-t')$ and activity $A_i$ at a given monomer $i$. The response matrix $\boldsymbol{J}$ of the polymer is given by Eq. (S76), as discussed in the previous section. In our linear model, we predict the following (discrete) mean squared separation map for a given profile of activity:

$$\boldsymbol{\Delta R}^2_{\text{pred}} = \sum_i \boldsymbol{S}_i\, A_i\,, \quad \text{where} \quad \boldsymbol{S}_i := \frac{\delta \boldsymbol{\Delta R}^2_{\text{pred}}}{\delta A_i}\,, \quad \text{[S77]}$$

which shows a linear but non-local response $\boldsymbol{S}_i$ to localized changes in activity. We now set up our optimization problem and seek the activity profile $A_i > 0\,\forall\, i$ that minimizes the mean squared deviation between the predicted mean squared separation map $\boldsymbol{\Delta R}^2_{\text{pred}}$ and a desired target conformation $\boldsymbol{\Delta R}^2_{\text{targ}}$:

$$F_{\text{fit}}(\{A_i\}) := \left\langle \boldsymbol{\Delta R}^2_{\text{pred}} - \boldsymbol{\Delta R}^2_{\text{targ}}, \boldsymbol{\Delta R}^2_{\text{pred}} - \boldsymbol{\Delta R}^2_{\text{targ}} \right\rangle_F\,, \quad \text{[S78]}$$

where $\langle \dots \rangle_F$ refers to the Frobenius inner product between two matrices[11]. By combining Eq. (S77) and Eq. (S78), we can write the *loss function* $F_{\text{fit}}(\{A_i\})$ as a quadratic equation,

$$F_{\text{fit}}(\{A_i\}) = \sum_{ij} A_i\, B_{ij}\, A_j - 2\sum_i V_i\, A_i + F_0\,, \quad \text{where} \quad \text{[S79a]}$$

$$B_{ij} = \left\langle \boldsymbol{S}_i, \boldsymbol{S}_j \right\rangle_F\,, \quad \text{[S79b]}$$

$$V_i = \left\langle \boldsymbol{S}_i, \boldsymbol{\Delta R}^2_{\text{targ}} \right\rangle_F\,, \quad \text{[S79c]}$$

$$F_0 = \left\langle \boldsymbol{\Delta R}^2_{\text{targ}}, \boldsymbol{\Delta R}^2_{\text{targ}} \right\rangle_F\,. \quad \text{[S79d]}$$

To minimize the *loss function*, Eq. (S79), given the constraint that the activity at each monomer is positive, $A_i > 0\,\forall\, i$, we use the Julia (9) library `Optim.jl` (10).

[9] We have here defined the lattice constant $a = L/N$ of the polymer, and related wave modes $q = n\pi/L$ to discrete wave coefficients $n$.

[10] That we need this term could be an artifact of having neglected effective translationally invariant correlations between the athermal excitations (i.e., a characteristic correlation length).

[11] For two matrices $\boldsymbol{Y}$ and $\boldsymbol{Z}$, the Frobenius inner product is given by the scalar $\langle \boldsymbol{Y}, \boldsymbol{Z}\rangle_F := \sum_{ij} Y_{ij} Z_{ij}$.

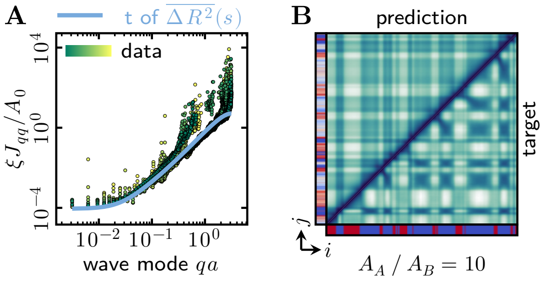

Fig. S8. **Reconstruction of activity profile based on data. A)** First, we marginalize the mean squared separation map $\Delta R^2(s, s')$ by averaging over diagonals with constant $s + s'$. We then use these data to fit the mechanical properties of the polymer, which are encoded in the response matrix. **B)** Comparison between the predicted mean squared separation map and our simulation data. The overall block structure matches well, but there are some features of our simulations that the linear model cannot capture. In the simulations, there are instances where two active polymer segments have a smaller spatial separation than their separation with an inactive segment that between them in sequence space. While the linear model can also show non-monotonicity in the mean squared separation as a function of genomic distance, it always predicts that active segments should increase their spatial separation.

### References

1. JG Kirkwood, J Riseman, The intrinsic viscosities and diffusion constants of flexible macromolecules in solution. *The J. Chem. Phys.* **16**, 565–573 (1948).
2. BH Zimm, Dynamics of polymer molecules in dilute solution: Viscoelasticity, flow birefringence and dielectric loss. *The J. Chem. Phys.* **24**, 269–278 (1956).
3. PE Rouse, A theory of the linear viscoelastic properties of dilute solutions of coiling polymers. *The J. Chem. Phys.* **21**, 1272–1280 (1953).
4. PC Parks, A. M. Lyapunov's stability theory—100 years on. *IMA J. Math. Control. Inf.* **9**, 275–303 (1992).
5. Z Gajić, MTJ Qureshi, *Lyapunov matrix equation in system stability and control*, Mathematics in sciences and engineering. (Academic Press, San Diego, CA), (1995).
6. K Kumar, On expanding the exponential. *J. Math. Phys.* **6**, 1928–1934 (1965).
7. O Hallatschek, E Frey, K Kroy, Tension dynamics in semiflexible polymers. *Phys. Rev. E* **75**, 031905 (2007).
8. H Zhang, et al., CTCF and transcription influence chromatin structure re-configuration after mitosis. *Nat. Commun.* **12**, 5157 (2021).
9. J Bezanson, A Edelman, S Karpinski, VB Shah, Julia: A fresh approach to numerical computing. *SIAM Rev.* **59**, 65–98 (2017).
10. PK Mogensen, AN Riseth, Optim: A mathematical optimization package for Julia. *J. Open Source Softw.* **3**, 615 (2018).
